# The genomic landscape of spider monkeys and northern muriquis from a conservation perspective

**DOI:** 10.1101/2025.03.07.641388

**Authors:** Núria Hermosilla-Albala, Marc Palmada-Flores, Jèssica Gómez-Garrido, Felipe Ennes Silva, Pol Alentorn-Moron, Armida Faella, Sira Martínez, Hugo Fernández-Bellon, Vanessa Almagro, Mariluce Messias, Mariane C. Kaizer, Izeni Farias, Tomas Hrbek, Maria N. F. da Silva, A. Patricia Mendoza, Fernando Vilchez-Delgado, Sam Shanee, José de Souza Silva Júnior, Rogerio Rossi, João Valsecchi, Pedro Mayor, Christina Hvilsom, Esther Lizano, Tyler S. Alioto, Marta Gut, Ivo G. Gut, Lukas F. Kuderna, Jeff Rogers, Kyle Kai-Hao Farh, Tomas Marques-Bonet, Jean P. Boubli

**Affiliations:** Institute of Evolutionary Biology (UPF-CSIC), Department of Medicine and Life Sciences, Universitat Pompeu Fabra. PRBB, C. Doctor Aiguader N88, 08003 Barcelona, Spain; Centro Nacional de Análisis Genómico (CNAG), C/Baldiri Reixac 4, 08028 Barcelona, Spain; Universitat de Barcelona, Barcelona, Spain; Research Unit of Evolutionary Biology and Ecology, Département de Biologie des Organismes, Université libre de Bruxelles (ULB), Brussels, Belgium; Mamirauá Institute for Sustainable Development, Tefé, Amazonas, Brazil; European Molecular Biology Laboratory (EMBL), Barcelona, Spain; Parc Zoològic de Barcelona, Parc de la Ciutadella s/n, 08003 Barcelona, Spain; Universidade Federal de Rondônia (UNIR), BR-364, Km 9.5, Sentido Acre, Porto Velho, RO, 76801-059, Brazil; Instituto Nacional da Mata Atlântica, Santa Teresa-ES, Brazil; Universidade Federal do Amazonas (UFAM), Manaus, Amazonas, Brazil; Instituto Nacional de Pesquisa da Amazônia (INPA), Manaus, Amazonas, Brazil; Neotropical Primate Conservation group, Department of Anthropology, Washington University, St. Louis, MO 63130, USA; Instituto de Medicina Tropical Alexander von Humboldt, Universidad Peruana Cayetano Heredia, Av. Honorio Delgado 430, San Martín de Porres, Perú; Cummings School of Veterinary Medicine, Tufts University, 200 Westboro Rd, North Grafton, MA 01536, USA; Neotropical Primate Conservation, Looe Hill, Seaton, Torpoint, Corwall, PL11 3JQ, United Kingdom; Museu Paraense Emilio Goeldi, Av. Gov Magalhães Barata, 376 - São Braz, Belém - PA, 66040-170, Brazil; Universidade Federal do Mato Grosso, Avenida Fernando Correa da Costa 2367, 78060-900, Mato Grosso, Brazil; Departament de Sanitat i Anatomia Animals, Facultat de Veterinària, Universitat Autònoma de Barcelona, Edifici V, Bellaterra, 08193, Barcelona, Spain; ComFauna, Comunidad de Manejo de Fauna Silvestre en la Amazonía y en Latinoamérica, Iquitos, Perú; Museo de Culturas Indígenas Amazónicas, Iquitos, Perú; Copenhagen Zoo, Roskildevej 38, 2000, Frederiksberg, Denmark; Institut Català de Paleontologia Miquel Crusafont (ICP-CERCA), Universitat Autònoma de Barcelona, Edifici ICTA-ICP, Cerdanyola del Vallès, Barcelona, Spain; Unidad de Paleobiología, ICP-CERCA, Unidad Asociada al CSIC por el IBE UPF-CSIC, Cerdanyola del Vallès, Barcelona, Spain; Illumina Artificial Intelligence Laboratory, Illumina Inc., Foster City, CA 94404, USA; Human Genome Sequencing Center and Department of Molecular and Human Genetics, Baylor College of Medicine, Houston, TX 77030 USA; Institució Catalana de Recerca i Estudis Avançats (ICREA) and Universitat Pompeu Fabra. Pg. Lluís Companys 23, 08010 Barcelona, Spain; School of Science, Engineering & Environment, University of Salford, Salford M5 4WT, UK

## Abstract

**Background:** Most populations of spider monkeys (*Ateles*) and muriquis (*Brachyteles*), two Neotropical primate genera, are under severe anthropogenic threats. Yet, taxon-wide population-level studies leveraging their degree of endangerment linked to their genetic diversity patterns and demographic history are lacking. To properly address this, there is a need to expand from morphological and genetic marker-based studies.

**Results:** We generated high-coverage genome sequencing for 58 individuals sampled across 8 *Atelidae* species, in the first population-wide study of all extant spider monkey species, in the wild and captivity, alongside northern muriquis (*Brachyteles hypoxanthus*). Additionally, we present a high-contiguity reference genome for *Ateles hybridus*. Here, we observe the overall levels of genetic diversity and genetic load of the analyzed populations do not align to their IUCN endangerment category. Moreover, we show that in the wild, genetic load is overall higher compared to the captive populations analyzed. Then, we depict two main trans and cis-Andean sister clades in *Ateles*, and further structure and dynamics outlined by the Madeira River in the latter clade. Lastly, we find that genes in highly divergent regions between *Ateles* and *B. hypoxanthus* are involved in central nervous system development and photorreception.

**Conclusions:** Our study shows i) the lack of concordance between the genetic diversity levels and extinction risk of these populations, suggestive of recent and strong external drivers; ii) increased genetic load in the wild in contrast to effective captive management, indicating mostly past demographic events; iii) structure and dynamics in spider monkeys that agrees with common biogeographical patterns and iv) genetic divergence between *Ateles* and *Brachyteles* potentially linked to distinct environmental light levels.

## Background

Spider monkeys (*Ateles*) and muriquis (*Brachyteles*) are two threatened genera of Neotropical primates belonging to the *Atelidae* family^1,2^ and include several Critically Endangered species. The distribution of the seven species in the *Ateles* genus (*A. belzebuth, A. chamek, A. fusciceps, A. geoffroyi, A. hybridus, A. marginatus* and *A. paniscus*) spans from Central America to South America, with species found both at the Pacific and Atlantic slopes^3^, mainly in lowland tropical forests, but sometimes also in high altitude^4^. Despite this, widespread deforestation and habitat degradation have led to reduced connectivity between *Ateles* populations, yielding considerable decreases throughout its range ^3,5^. On the other hand, *Brachyteles hypoxanthus* is found in protected areas of the diverse Brazilian Atlantic Forest in the states of Minas Gerais, Espírito Santo and Bahia^2^. The habitat of *B. hypoxanthus’* populations is currently severely fragmented^6,7^, with an estimated census of less than 1000 individuals^2^ thus highly vulnerable to genetic erosion^6^. In spite of the overall large distribution range covered by these atelid species, they face anthropogenic threats consisting of habitat destruction and fragmentation due to deforestation, hunting and poaching^2,5,8–15^. Moreover, spider monkeys and northern muriquis share unique characteristics that make them especially susceptible to the aforementioned challenges: They have extended life histories with exceptionally long interbirth intervals when compared to catarrhine primates of similar size^6,15^. This is longest in *A. belzebuth* (43.7 ± SD 5.1 months)^16,17^, in contrast to 22 and 24 months on average in macaques and baboons respectively^18^. Additionally, spider monkeys require very specific ecological conditions due to their specialized frugivorous diet^3,5,19^.

Maintaining the genetic diversity of a species to ensure its long-term survival is a key factor in conservation genomics^20^. This further includes the maintenance of low levels of inbreeding genome-wide and reduced load of detrimental mutations. Detailed conservation strategies are nowadays directed to both *in-situ* and *ex-situ* populations^21^. *Ex-situ* conservation programmes are tailored to each species, as those for *A. fusciceps, A. hybridus* and *A. paniscus* led by the European Association of Zoos and Aquaria (EAZA)^22^. On the other hand, the direct consequences of anthropogenic threats on *in-situ* populations are oftentimes overlooked due to their multilayered complexity. However, they can lead to population decline through fragmentation, thus intensifying the effect of genetic drift in the long run. In addition, the impact of environmental threats on population survival can be intensified by the demographic past of these. Accordingly, the study of endangered species from an evolutionary population genomics perspective, including their levels of genetic diversity, inbreeding, genetic load, and dynamics, can be widely informative to conservation strategies while in full consideration of current external threats.

However, studies including the whole set of *Ateles* species are scarce and based either on morphological traits such as coat color^23^, karyotypic data^24^, or on a reduced number of genetic markers (e.g. RFLPs^25^, mitochondrial genes^1,4^, ultra conserved elements (UCEs)^26^). Furthermore, given their intricate evolutionary trajectories, the classification of spider monkeys has been revisited multiple times without great consensus, as has been the case in many platyrrhine taxa^27,28^. In addition, while recent studies leveraging genetic data explore the population dynamics of individual *Ateles* species^16,29^ are informative for conservation efforts, these provide limited context for a broader taxon-wide view. Through the latter, potentially meaningful interspecies interactions and shared patterns could be identified, such as common barriers to dispersal or the effect of ecological features.

Here, we sequenced the whole genomes of 58 individuals from 8 Atelidae species sampled from 9 populations to evaluate the genome-wide heterozygosity as a proxy of genetic diversity, inbreeding and genetic load of spider monkeys and northern muriquis.

We present the first population-wide study of spider monkeys based on high-coverage whole genome sequencing data of all extant species from both captive and wild individuals, together with northern muriquis. We furthermore introduce an *Ateles hybridus* high-contiguity reference genome assembly generated as part of two greater efforts to study and preserve Critically Endangered species: ORG.one^30^ and Cryozoo^31^. We examine the genetic makeup of all *Ateles* species and *Brachyteles hypoxanthus* with the aim to describe their levels of genetic load, diversity and inbreeding in captive and wild populations from an evolutionary perspective. We provide a detailed description of these populations’ structure and phylogenetic relationship, by focusing on a genome-wide set of variants for more than one individual in almost all species. Consistent with previous observations in many other taxa across the Amazon, we illustrate how rivers^27,32,33^ but also current distribution ranges outline the genetic structure of spider monkeys, and additionally determine the dynamics of some of the *in-situ* populations analyzed. Lastly, we also explore the genomic regions underlying the differentiation between *Brachyteles hypoxanthus* and the *Ateles* genus in an effort to describe their evolutionary trajectories after their split ca. 11 Mya^34^.

## Results

We sequenced the genomes of 58 unrelated individuals to a median coverage of 18.3x and a range from 5.5x to 34.6x, and identified 168.096.572 single nucleotide polymorphisms (SNPs) after applying hard filters. 15 and 43 samples in our dataset were derived from individuals of captive and wild origin respectively, covering all species in the *Ateles* genus and *Brachyteles hypoxanthus*. Based on the geographic location where *A. chamek* individuals were sampled relative to the Madeira River and its right hand tributary, Guaporé River, these were further divided into two independent populations following hypotheses based on prior observations^5^: *A. chamek_NW* (northwestern) and *A. chamek_SE* (southeastern), previously referred to as *A. longimembris*^35^ (Fig. 1A). The number of individuals per sampled population was highly uneven particularly for wild ones given the difficulty of obtaining biomaterials, ranging from 1 to 19 in wild populations and from 2 to 13 in captive (Supplementary Data 1).

**Figure 1.**
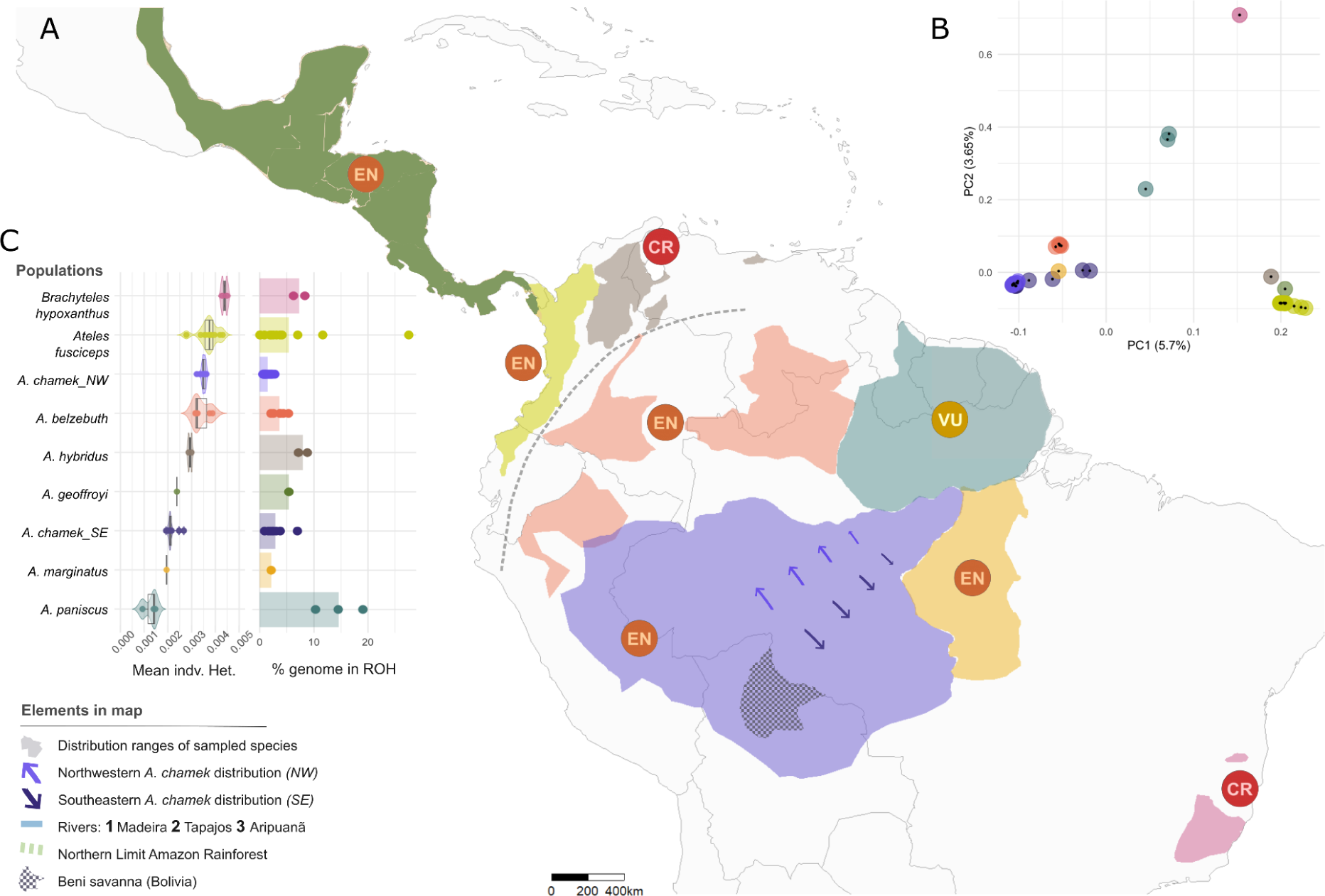
Spider monkeys and northern muriquis populations. A) Distribution ranges with relevant geographic features of sampled species together with their endangerment status based on the IUCN Red list of Endangered Species: CR - Critically Endangered, EN - Endangered, VU - Vulnerable; B) Principal component analysis of unrelated samples based on 58.20M SNPs. Percentages denote variance explained by each PC; C) Mean individual heterozygosity distribution and percentage of the genome in ROH per population.

### IUCN conservation status does not correspond with spider monkeys’ nor northern muriquis’ genetic makeup

The mean population heterozygosity in the callable genomes of spider monkeys ranged from 0. 0013 to 0. 0040 and was 0. 0044 in northern muriquis (Fig. 1C), being overall higher than previous observations in atelids^26^.

After *B. hypoxanthus,* with the highest heterozygosity genome-wide (*Het*. = 0. 0044), we find the mostly *ex-situ* sampled *A. fusciceps* (*Het*. = 0. 0038), *A. chamek_NW* (*Het*. = 0. 0035) and *A. belzebuth* (*Het*. = 0. 0034) presented close heterozygosity values yet distinct genome-wide distributions (Fig. 1C, S10-S12). On the contrary, *A. paniscus* (*Het*. = 0. 0013) was by far the least genetically diverse population, preceded by *A. marginatus* (*Het*. = 0. 0019) and *A. chamek_SE* (*Het*. = 0. 0021) (Fig. 1C).

Congruently presenting the lowest genome-wide heterozygosity, *A. paniscus* had the largest number of short (0.5-1Mb) and intermediate size (1-2Mb) runs of homozygosity (ROHs) when compared to the other populations (Fig. 2A). While *B. hypoxanthus* was at the opposite extreme in terms of genome-wide heterozygosity, it presented the largest number of short ROHs in the dataset after *A. paniscus* (Fig. 1C, 2A). Contrary to long ROHs, which are indicative of recent inbreeding, short ones typically point to an older demographic event, the signal of which has been broken down by recombination through generations^36^. Therefore, the observed amount of short homozygous tracts could be suggestive of either an old population bottleneck or the long-term persistence of population structure in the past followed by fragmentation, as proposed by Chaves *et al.* (2011)^6^.

**Figure 2.**
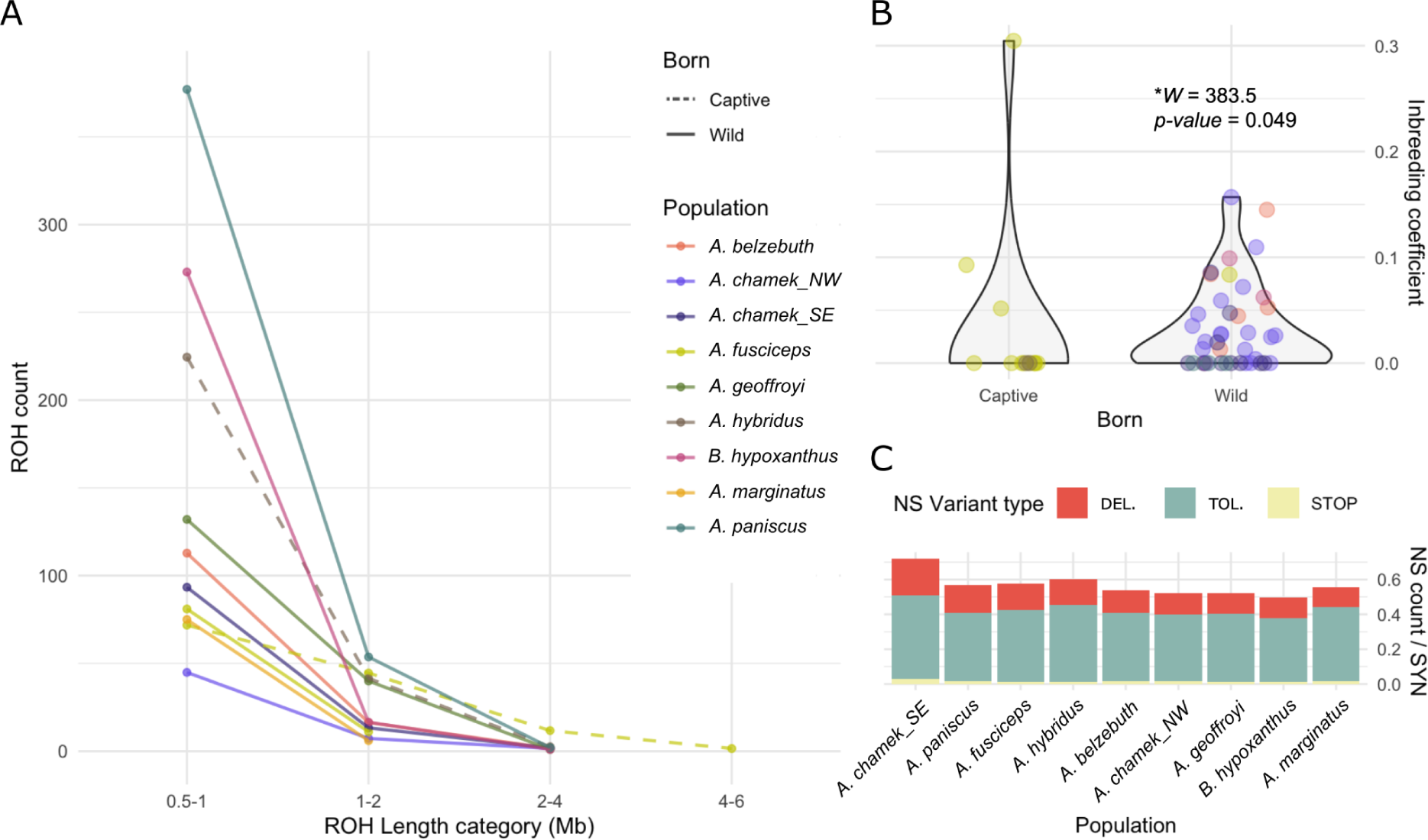
Inbreeding and mutational load in *Ateles* and *Brachyteles hypoxanthus*. A) Count of runs of homozygosity (ROHs) above 0.5Mb by length category and sampled population as well as management status. B) Genome-wide inbreeding coefficients in captive vs. wild individuals coloured by population. Statistic and significance of non-parametric Wilcoxon test indicated. C) Ratio of non-synonymous (NS) mutations divided by number of synonymous (SYN) in each population. Coloured by NS mutation type as predicted by SIFT: deleterious (DEL.), tolerated (TOL.) and STOP.

With contrasting phenotypes, distinct forest type preferences and an estimated divergence time of ca. 11 Mya^34^, spider monkeys and northern muriquis showed marked genetic dissimilarities (Fig. 1A-C, S1). Consistent with previous studies^20,26^, none of the analyzed parameters linked to the genetic diversity, inbreeding nor genetic load of the studied populations corresponded to their assigned IUCN Red List conservation status (Fig. 1A,C) in terms of susceptibility. This likely indicates the consequences of the generally great extinction risk of these species are overall not strongly reflected in their genomes yet ^37^. For example, *A. hybridus,* as a Critically Endangered (CR)^12^ species, exhibited intermediate values in the genus of the latter parameters (Fig. 1C, S8). Nevertheless, we also observe the opposite case, where *A. paniscus* presents the lowest threatened category of all spider monkey species (Vulnerable - VU)^14^ while showing the highest proportion of the genome in ROH and lowest genome-wide heterozygosity (Fig. 1C, S8).

### Overall inbreeding and genetic load are not more prevalent in captive populations compared to wild ones

Although large variation was found among *A. fusciceps* captive individuals, some of these were unique in displaying ROHs longer than 4Mb, suggesting recent inbreeding contrary to the only wild sampled individual in the species (Fig. 2A). Furthermore, captive *A. fusciceps* individuals showed the largest proportions of the genome in homozygosity (fROH), although this was also observed in wild sampled *A. paniscus*. fROH values in these samples ranged from 0. 02% to 27. 54% and from 10. 31% to 19. 10% respectively (Fig. 1C, S8, S14).

In spite of captive *A. fusciceps* PD_1568 exhibiting an outlier inbreeding coefficient (IC) above 0. 3 (Supplementary Data 1 – Inbreeding_coeffiicient column, Fig. S13), the ICs in captive (*Range* = [0 − 0. 09], without outlier) and wild (*Range* = [0 − 0. 16]) individuals were nominally significantly different (*W* = 383. 5, *p* − *value* = 0. 049), overall being more prevalent and higher in wild populations (Fig. 2B). Notably, *A. chamek_NW, A. belzebuth* and *B. hypoxanthus* presented widespread inbreeding (Fig. 2B), yet showed some of the highest mean genome-wide heterozygosity values (Fig. 1C).

Likewise, there was no concordance between the captive status of the sampled individuals and their observed genetic load. The latter was approximated by the proportion of non-synonymous deleterious mutations normalized by the observed synonymous mutations 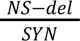. For this, the wild *A. chamek_SE* population showed the highest average ratio (Fig. 2C). This was followed by *A. paniscus* and *A. fusciceps,* which presented very similar proportions, whereas *A. marginatus* showed the lowest genetic load proportions (Fig. 2C).

Along with *A. chamek_SE, A. belzebuth* displayed significantly different values of 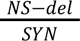 inside and outside of ROHs (Fig. S16). However, opposite to *A. chamek_SE, A. belzebuth* exhibited a significant enrichment of detrimental mutations inside ROHs, specifically in short ones (Fig. S17). This could potentially be linked to this species’ extremely long interbirth intervals while found in a patchy habitat^16^. The latter was also the case only in *A. hybridus*, however the enrichment in ROHs was not significant (Fig. S16).

### The Andes mountain range and the Madeira River outline the genetic structure of spider monkeys

The genetic structure in the *Ateles* genus is highly concordant with the geographic distribution of the enclosed populations (Fig. 1A, B): Those found in Central and Northern South America (trans-Andean group: *A. geoffroyi, A. hybridu*s, *A. fusciceps*) formed an independent genetic cluster from those below the Andean range at the northern limit of the Amazon rainforest (cis-Andean group: *A. chamek_SE, A. chamek_NW*, *A. marginatus*, *A. belzebuth*) based on PC1 (5.79%) and the genome-wide phylogeny (Fig. 1A, S1, S2). The observed structure outlined by the Andes is a common biogeographical pattern, where species on the west of these in Central America (trans) are usually the sister clade to species on the east in the Amazon (cis)^38^. Nonetheless, even though *A. paniscus* clustered with the cis-Andean group in accordance with its own geographic distribution based on these analyses, it was the most differentiated population in the genus based on PC2 (3.56%) (Fig. 1A, S2). The observed ranges of genetic diversity and inbreeding (Fig. 1B, C, 2) were not distributed in alignment with the described population structure nor with the phenotypic characteristics of each species in terms of coat patterns^3^.

When *B. hypoxanthus* was included, this species separated from all *Ateles* populations based on ADMIXTURE (Fig. 1A, B, S6) and PCA, with the trans-Andean cluster closest to it based on PC1 (5.7%) and *A. paniscus* based on PC2 (3.65%) (Fig. 1B). Furthermore, the two populations of *A. chamek* described here with respect to the Madeira River exhibited corresponding signals of genetic distance. These were differentiated in the PCA analyses (Fig. 1B, S2, S3), exhibited distinct genome-wide heterozygosity distributions (Fig. S9), were soundly discriminated by ADMIXTURE at best *K* = 5 (Fig. S6) and by phylogenetic clustering with high support (*Phylogentic_Support* = 0. 97) (Fig. S1).

**Figure 3.**
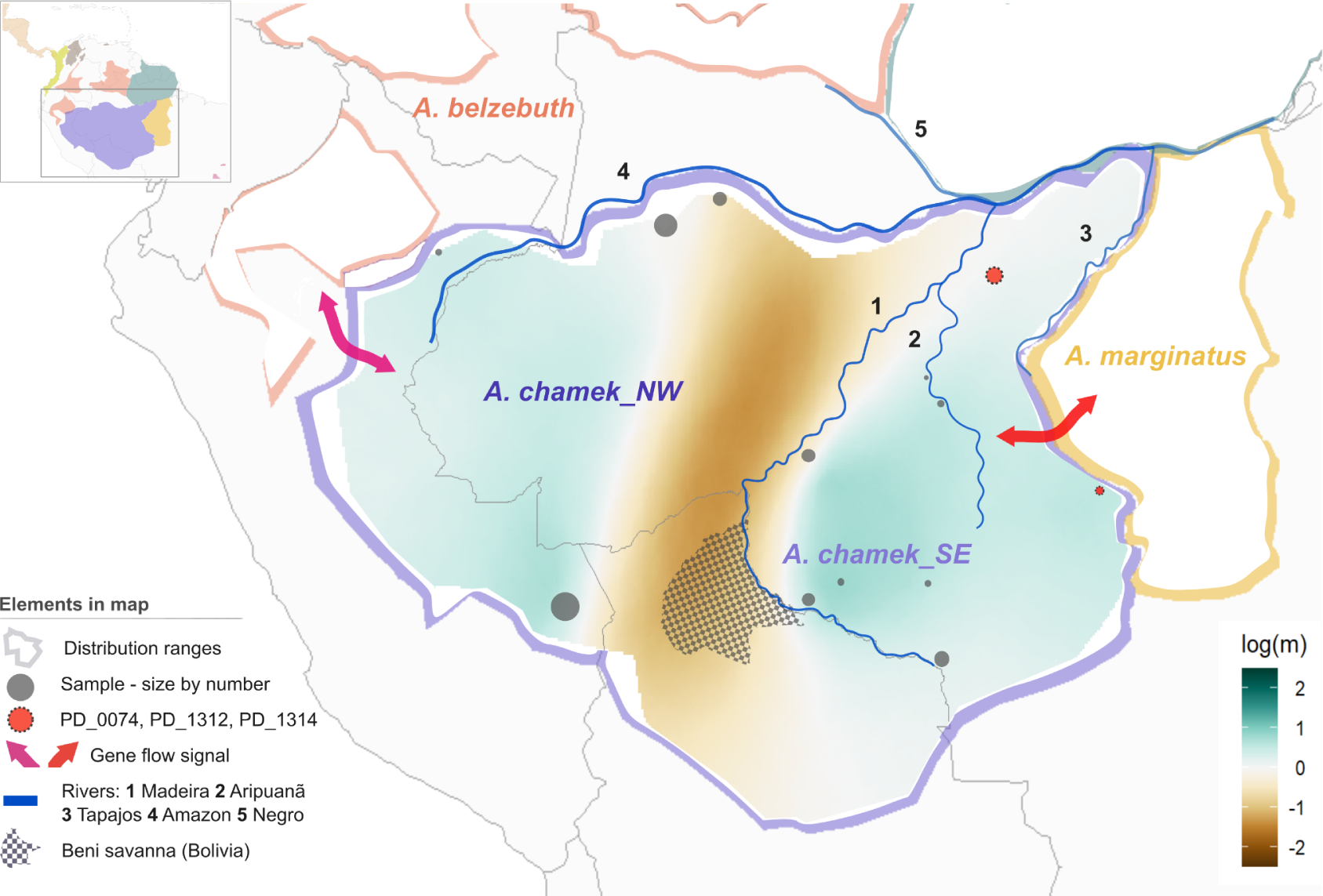
Population dynamics of Amazonian spider monkeys. Estimated effective migration surfaces (EEMS) in the distribution range of *A. chamek* depicting the Madeira River’s course and the Beni savanna as barriers to migration isolating northwestern and southeastern parts of its distribution. Indicated significant gene flow signals detected at opposite banks of the Madeira River between independent *A. chamek* populations and *A. belzebuth* and *A. marginatus* respectively. *A. chamek_SE* samples driving the latter signal highlighted.

The genetic closeness to *A. marginatus* observed in *A. chamek_SE* in contrast to *A. chamek_NW* in terms of genetic structure (Fig. 1B, S1, S2) was further tested and yielded a significant gene flow signal only between the first two (*f* − *branch* = 0. 075) (Fig. 3, S18). In agreement with our hypothesis, the aforementioned observations could be explained by the discontinuity to the distribution of *A. chamek* through its evolutionary history caused by the Madeira River. This was further supported by the observed deviations from isolation by distance in terms of reduced migration coinciding with its course in the habitat range of the species (Fig. 3). This signal also overlapped with the Beni savanna in Bolivia towards its headwaters, pointing to further allopatric barriers through the river course at both extremes of the species’ distribution (Fig. 3).

Moreover, after a comprehensive examination of the population assignment of the sampled individuals, we observed further substructure and stronger genetic proximity to *A. marginatus* in a few *A. chamek_SE* individuals (PD_0074, PD_1312, PD_1314) in contrast to the rest of the population. These three individuals were closer to *A. marginatus* in the PCA of cis-Andean populations (Fig. S3), the phylogenetic tree (Fig. S1), and shared totally or partially its ancestry component at *K* = 8 (Fig. S7). Although the latter *K* exhibits a high cross-validation error, it is reported as a reflection of population structure rather than actual ancestry sharing. Moreover, these three individuals were sampled in a region where the Aripuanã River divides the inter fluvial area between the Madeira and Tapajos rivers from the other *A. chamek_SE* sampled locations (Fig. 3). There, the Aripuanã River is known to delimit the distribution of other primates such as *Plecturocebus miltoni*^39^, yet, this potential barrier is not captured by EEMS. Consistent with the potential isolation experienced by these individuals’ population, PD_1312 and PD_1314, with the same sampling coordinates (Fig. 3), presented the highest proportion of 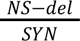 in all the dataset but low or values of fROH (Fig. 1C, S8, S19). Furthermore, these three individuals yielded an *f-branch* of 0. 036 with *A. marginatus* when considered as an independent population from the other *A. chamek_SE*, which showed no gene flow signal here (Fig. S19), indicating the initial signal identified (Fig. S18) was most likely driven by this subpopulation.

Lastly, consistently with both populations being distributed across the northern bank of the Madeira River, significant *f*-branch values were identified between *A. belzebuth* and *A. chamek*. Specifically, this was 0. 05 and 0. 06 respectively between *A. belzebuth* and *A. chamek_NW* and between the first and the ancestor of both *A. chamek* populations (Fig. 3, S18). The last could be explained by the proposed hypothesis that this species originated in the northern part of their distribution^1^.

### Genomic regions driving the divergence between spider monkeys and northern muriquis encompass genes involved in photoreception and the MAPK/ERK pathway

Regions in the genome with an *Fst* index found in the 97th percentile in each *Ateles* population to *B. hypoxanthus* comparisons overlapped in 307.141 base pairs which spanned over 6.886 positions in 8 genes: *BRAF, CDC42BPB, FNDC3A, GNAT1, ITPR2, MARK3, RBM6* and *WDPCP*.

Focusing on the functional enrichment of these genes, firstly based on the Molecular Signatures Database^40^, four of these (*BRAF, ITPR2, MARK3* and *RBM6*) presented the PU1_Q6 motif that acts as binding site to *SPI1* transcription factor, which could be indicative of coevolution. This is involved in the innate immune response^41^ and nervous system development^42^. Secondly, *BRAF* and *ITPR2* drove the enrichment of synaptic activity-related pathways *Long-term depression* and *Long-term activation* pathways when using FUMA^43^ in addition to *Parathyroid hormone synthesis, secretion and action* and *Serotonergic synapse* when using WebGestalt^44^ (*FDR* < 0. 05) in the KEGG PATHWAY Database^45^. Lastly, we also used WebGestalt to examine enrichments in the REACTOME Database^46^, where *BRAF* and *MARK3* drove the identification of the following categories related to the MAPK/ERK cell signaling pathway (*FDR* < 0. 05): *Signaling by high-kinase activity BRAF mutants, RAF activation, MAP2K and MAPK activation, Signaling by RAF1 mutants, Negative regulation of MAPK pathway, Signaling my moderate kinase activity BRAF mutants, Signaling by RAS mutants, Paradoxical activation of RAF signaling by kinase inactive BRAF, Signaling downstream of RAS mutants, Signaling by BRAF and RAF1 fusions.* For each database, significant categories were reported in increasing enrichment ratio.

On the other hand, although not found in any enriched category, other identified genes can be highlighted with functions related to photoreception: while *GNAT1*^47^ is active in rods thus critically involved in light vision, mutations in *WDPCP* have been implicated in cone-rod dystrophy through photoreceptor disruption in the Bardet Biedl Syndrome^48^ and mutations in *CDC42BPB* related to retinal degeneration^49^.

## Discussion

The investigation of population-level patterns in spider monkeys and northern muriquis populations from a genome-wide perspective for the first time, offers a groundbreaking opportunity. Alongside with a new reference genome, this allows us to more deeply understand the actual degree of threat to these species linked to their genetic diversity patterns. Inherently tied to this, a detailed description of their respective evolutionary relationships provides an overview of the patterns that have shaped their genetic makeup. All this becomes particularly relevant in light of the degradation and fragmentation South American tropical forests are experiencing, in particular the periphery of the Amazon rainforest and the Braziliian Atlantic Forest^2,50^, which are the habitats of the studied populations. Widespread deforestation linked to human activity has put some of these species among the most threatened primates on earth^2,51^. Hunting and land conversion for agriculture, logging, mineral and fossil fuel mining are destroying and transforming their habitat into shrinking fragmented forest patches, as has been observed in the diminished actual area of occupancy of spider monkeys in their original distribution range^5,6,52^. Additionally, the life-history traits and ecological properties of all studied populations, such as long interbirth intervals causing delayed population turnover^16^, their preference for the canopy layer in the forests^5,53^, as well as spider monkeys-specific ranging behaviour shaped by the availability of fruits^3^ whilst exhibiting high body mass, make them particularly vulnerable to anthropogenic threats. Contrary to northern muriquis, which may prefer to feed on leaves^54^ hence not being so limited by preferred resources, spider monkeys are unable to survive in extremely patchy and fragmented, second growth forest habitats^55,56^.

The importance to preserve these species must be underlined by the profoundly significant seed disperser role spider monkeys play in the ecosystems of neotropical forests throughout their broad range^5,15^. Disruptions on these habitats negatively affecting the survival of spider monkeys would retroactively translate in paramount alterations in these biomes, which are in turn vital for the earth’s natural stability.

Due to the difficulty of obtaining biomaterials from wild populations, our dataset is limited by an uneven representation of the studied populations in terms of sample number and sequencing coverage. We understand that this may bias the outcome of some analyses, for example by underestimating the heterozygosity of some of the populations, in particular that of *A. geoffroyi* and *A. marginatus* (Fig. 1C), given that population-level analyses prompt variant discovery. The latter is well exemplified by comparing the genetic diversity values reported here in comparison to those in Kuderna *et al.* (2023)^26^ for most of the spider monkey species. Observed values are overall higher in this study in contrast to the latter, where one individual was analysed while applying comparable approaches. Furthermore, though we are able to draw relevant conclusions by comparing wild and captive populations, we are unable to make a thorough comparison of these in any given species. This as well, comes as a consequence of the lack of an even number of samples of each origin in all species.

### 1. Structure, dynamics and genetic load in the wild

Previous descriptions of the evolutionary relationships among spider monkey species have not found a consensus based on their karyotypes^24^, reduced representations of their genomes^25,26^, gene trees^1,4^ nor morphological traits^57^, as these likely fail to represent the complete history of these taxons. Here, using whole callable genomes of all species, we estimate their population structure (Fig. 1B, S2-4, S6-7) and phylogeny (Fig. S1), which we found most similar to the clustering outlined by ultraconserved elements in the genomes of one single individual for some of the species analyzed here^26^. Moreover, our observations agreed with the other approaches in that the pairs of species *Ateles fusciceps* and *A. geoffroyi,* and *A. chamek* and *A. belzebuth* respectively shared the most recent common ancestors in the clade. In line with this, we describe how the geographic distribution of the *Ateles* species aligns with the genetic proximity of these when *B. hypoxanthus* is used as an outgroup. In this, the Andean mountain range at the northern limit of the Amazon rainforest distinguishes the two main sister clades in the genus (Fig. 1A, B). Lastly, in the Amazonian distribution of spider monkeys, the Madeira River also plays a crucial role in delimiting *A. belzebuth, A. chamek* and *A. marginatus’* population dynamics (Fig. 3).

The large area inhabited by spider monkeys encloses a myriad of distinct past scenarios, contemplating also that of northern muriquis, some areas have suffered greater changes since the last glacial maximum (LGM), including changes in Amazonian river drainage networks^58^, forest shrinkage and expansion due to cyclic environmental changes, and anthropogenic forest fragmentation. However, other geographical features and environments have been more stable, for example western to central Amazon, where wet tropical forest has persisted ^59^. Altogether, this variability has facilitated divergent and distinct evolutionary trajectories as observed in other studies on platyrrhine species^27^. As an overall trend, we observe how current events may have not yet left a strong footprint in the genomes of some of these species that we have been able to detect. However, we found that these signals were still outlined by past demography. In most cases, this resulted in a lack of correspondence between the assigned IUCN category of the species and their genetic makeup^2,8–14^ (Fig. 1A, C, 2A-C). For example, we observed that *B. hypoxanthus* exhibits high levels of genome-wide heterozygosity in comparison to the reported values for all *Ateles*^26^ and low genetic load in spite of its Critically Endangered status (Fig. 1C, 2A, C). The latter species nonetheless, presented a large number of short runs of homozygosity, which due to their length can be linked to past demographic events. Yet, given our sampling limitations, we were not able to fully decipher whether this pattern reflected a past population bottleneck or long-term population structure and fragmentation, which Chaves *et al.* (2011)^6^ propose.

#### 1.1. *Ateles’* Amazonian populations

On the other hand, the distribution of genetic diversity, mutational load and inbreeding in spider monkeys does align with the geographic distribution of the species for the most part, with some exceptions (Fig. 1A, C, 2A-C). Focusing on Amazonian *Ateles,* it has been reported that in particular eastern, but also southern regions of the Amazon rainforest underwent great transformations during the LGM. These areas likely went through periods of drier conditions leading to the expansion of savanna-like environments and thus becoming unavailable habitats for *Ateles* species^58,59^. In accordance with a potential past population bottleneck related to habitat instability, we observe how *A. paniscus,* found in the easternmost distribution of the genus in the Guiana shield^60^, exhibits the lowest genome-wide heterozygosity. In this, we also observed elevated genetic load: the greatest number of short ROHs, fROH and the second highest proportion of deleterious mutations in the dataset (Fig. 1A, C, 2A, C). Likewise, distributed through the southeastern part of the Amazon rainforest, *A. marginatus* presents the second lowest genome-wide mean heterozygosity in the genus (Fig. 1A, C). *A. paniscus* and *A. marginatus* have been identified as potentially the most severely impacted species if the extreme deforestation predictions for the Amazon are realized by 2050^50^. Moreover, the latter is already found across the current Amazonian arc of deforestation^1,61^. In terms of population susceptibility, we underscore the misalignment of this biogeographical information, the observed patterns of genetic diversity, inbreeding and genetic load and their current consideration as Vulnerable (VU)^14^ and Endangered (EN)^13^ respectively by the IUCN (Fig. 1A).

In other parts of Amazonian *Ateles’* distribution, we find that the presence of major geographic barriers to dispersal, as the Madeira River, do align with the distribution of genetic diversity (Fig. 3), agreeing with the long standing paradigm of the riverine barrier hypothesis ^32^. As such, *A. chamek_SE* and *A. marginatus*, found at the southern bank of this, presented diminished heterozygosity, increased genetic load and short ROHs in the case of *A. chamek_SE.* This increase is in comparison to *A. belzebuth* and *A. chamek_NW*, that present a slight habitat overlap at the opposite bank of the river (Fig. 1A, C, 2A, C). However, we observed widespread genome wide inbreeding in the individuals of the last two populations (Fig. 2B). This was surprising given these exhibited the highest levels of genome-wide mean heterozygosity in the wild, probably associated with larger and less perturbed ancestral populations. Moreover, *A. belzebuth* showed a significant enrichment of deleterious mutations in short homozygous tracts (Fig. S17). Altogether, these results could be indicative of increased drift resulting in the inability to purge slightly deleterious mutations through purifying selection. A plausible explanation for this could be the exceptionally long interbirth intervals observed in this species, the longest within the genus^16^, combined with its habitat being in a very patchy distribution range^8^ (Fig. 1A).

Through its imposition as an effective allopatric barrier, the dynamics of the four aforementioned populations are greatly governed by the Madeira River (Fig. 3). This has also been proposed for other primate species such as titi monkeys (*Plecturocebus spp.*)^62,63^. In light of all these observations, we support Grant (2023)^5^ in highlighting the independence of the two studied *A. chamek* populations to properly tackle their preservation. Accordingly, we identified distinct historical migration patterns in these two. First, significant gene flow was observed between *A. belzebuth* and *A. chamek_NW* (Fig. 3, S18). We hypothesize this could take place towards the westernmost part of their distribution and the headwaters of the Amazon River (Fig. 3), since the latter could represent a barrier to dispersal otherwise. Moreover, we found that *A. belzebuth* also exhibited significant gene flow signals with the ancestor of both *A. chamek* populations (Fig. S18). This could be linked to *A. belzebuth* and *A. chamek_NW* exhibiting signs of potentially larger ancestral effective population sizes. This last identified signal would agree with i) the theory that *A. chamek* originated in the northwestern area of its distribution^1^; and with ii) the observed signs of strengthened genetic drift in *A. chamek_SE,* similarly to mexican populations of *Alouatta palliata*^64^, as a reflection of an ancestral founder population (Fig. 3, 2A, C). Secondly, even though *A. chamek_SE* as a whole exhibited signs of gene flow with *A. marginatus* (Fig. 3, S18), this signal was driven by a set of three individuals (PD_0074, PD_1312, PD_1314). They were sampled in an area where the Aripuanã River divides the interfluve between the Madeira and Tapajos rivers (Fig. 3, S19). The latter observation would not necessarily rule out the possibility that there has been connectivity between *A. marginatus* and the remaining *A. chamek_SE* individuals. Nevertheless, it implies that the signal driven by PD_0074, PD_1312 and PD_1314 is strong enough to overshadow what could be detected in the other *A. chamek_SE* individuals. The high levels of deleterious variation (Fig. S19) and population structure (Fig. S1, S3, S7) exhibited by PD_0074, PD_1312 and PD_1314 suggest they could belong to an isolated population. This would hypothetically be isolated from the remaining *A. chamek_SE* individuals by the Aripuanã River, which has previously been described to be an effective barrier to dispersal for other primates such as *Plecturocebus miltoni* populations^39^. Yet, this potential barrier did not apparently cause deviations from isolation by distance (Fig. 3), which could be suggestive of some alternative scenarios. These encompass the possibilities that i) this population may have been historically isolated but has not been for a while, or ii) that other undetected non-allopatric barriers are at stake.

### 2. Captive and wild spider monkeys

Retaining the genetic diversity of captive populations is a key goal for *ex-situ* modern breeding programs as a proxy for the maintenance of their fitness potential^65^. This is observed in the captive sampled *A. fusciceps* in our dataset: These exhibited the highest values of heterozygosity in all examined spider monkeys while mainly composed by captive individuals (Fig. 1C). On the other hand, though this comparison is broadly limited by sample count, wild *A. fusciceps* PD_2714 was the exception to the species’ high diversity, found at the lower tail of the species’ distribution. Moreover, the two *A. hybridus* captive individuals analyzed also exhibited higher genome-wide mean heterozygosity than the average in the genus. In contrast, in spite of being part of the trans-Andean clade together with the former two and presenting an actually larger distribution range in the wild than them^5,10–12^, the wild sampled *A. geoffroyi* did not follow their genetic diversity patterns (Fig. 1A, C).

*Ex-situ* conservation programs nonetheless present intrinsic limitations. Since they focus on a reduced representation of each of the managed species, these work on a limited number of individuals and genetic pool. All this makes these populations more susceptible to genetic drift and inbreeding^65^, through departing from panmictic mating and reducing the possibility of gene flow.

In the aforementioned species, some of the *A. fusciceps* captive individuals present the longest ROHs in the dataset, as well as the highest number of intermediate-size ROHs (Fig. 2A). This is indicative of recent inbreeding likely linked to their captive status in contrast to the single wild individual of the species (Fig. 2A). Yet, great variation is found in the set of captive managed *A. fusciceps* individuals (Fig. 1C, S8), which could be attributed to the number of generations the analyzed lineages have been bred in captivity^36^. Similarly, *A. hybridus* lacks long ROHs while it presents an equally elevated count of intermediate sized ones to wild *A. geoffroyi* and captive *A. fusciceps.* Moreover, it exhibits an even higher number of short tracts of homozygosity reflected in an fROH of more than 15% (Fig. 1C, 2A). Nevertheless, the observations in *A. hybridus* likely reflect past demographic events rather than the effects of *ex-situ* management^36^.

Then, we find that genome-wide inbreeding coefficients in captive populations fall inside the distribution of wild individuals’ with the only exception of *A. fusciceps* PD_1568. Actually, genome-wide inbreeding is more widespread and nominally significantly higher in wild populations (Fig. 2B). This result may speak of the overall effectiveness of management programs in these species, while giving a warning on the resilience potential of wild populations linked to past demographic events^65^. In line with this, wild *A. paniscus* and *A. chamek_SE,* driven by PD_0074, PD_1312 and PD_1314, exhibit higher genetic load than captive *A. fusciceps* and *A. hybridus* on average (Fig. 2C). Relevantly, the only wild *A. fusciceps* is the 10th sample with the highest deleteriousness ratio in the dataset (Fig. S19).

### 3. Evolutionary trajectories of *B. hypoxanthus* and *Ateles*

*B. hypoxanthus* and *Ateles* are thought to have diverged ca. 11 Mya^34^. This divergence is notably visible in terms of genetic makeup (Fig. 1A-C, S1) and phenotypic traits, given the first are of larger body size and lighter color coats^66^. But also ecologically, with distinct dietary and habitat preferences: Spider monkeys inhabit dense tropical forests and mostly depend on soft fruits^3,56^. On the contrary, northern muriquis are found in the Brazilian Atlantic Forest, where the canopy is more fragmented due to the rugged relief and steep mountainous terrain and mostly feed on leaves^2,54^. Although both taxons are mostly found in the canopy layer^5,53^ of their distinctly preferred forest types, the distinct topography, tree species and growth patterns cause crucial differences in the sparseness of the respective canopies. In lowland tropical forests inhabited by *Ateles* species, we find a densely layered canopy^5^. Contrarily, this is lower, more open, fragmented^53^ and therefore brighter in Atlantic Forest’s hills and mountains. We hypothesize that the amount of light in the respective habitats of spider monkeys and northern muriquis together with the exposed differences in feeding behaviours and diet restrictiveness, could explain the identification of *GNAT1*^47^*, WDPCP*^48^ and *CDC42BPB*^49^ genes, which are involved in photoreception, in genomic regions of high genetic differentiation between the two groups.

Furthermore, we observed the associated genetic dissimilarities to their divergence were overrepresented by regions in the respective genomes associated with the *MAPK/ERK* pathway. This plays a pivotal role in cell cycle progression, survival and differentiation^67^. Also, it has recently been linked to the development of social and emotional behaviors through the mechanisms it mediates in the amigdala^67^, as well as to neuronal and synaptic plasticity in humans^68^. Accordingly, the same genes identified as part of this pathway were enriched for synaptic activity-related functions. In parallel, we found that the identified genes in regions of high divergence were alternatively or additionally targets of the *SPI1* transcription factor, involved as well in the innate immune response^41^ and the development of the nervous system^42^. Together, all these observations could indicate that central nervous system development has been relevant in the divergence of these two taxa.

## Methods

### *Ateles hybridus* genome assembly

#### Sample collection and processing

We expanded fibroblast cell lines from the CryoZoo biobank for individual 12393 from The Barcelona Zoo in T75 flasks. When the cells reached confluency, we rinsed with PBS, harvested these and added 5 mL of 0.05% Trypsin-EDTA. We then incubated the flasks at 37°C for 4-5 minutes to detach the cells. Once detached, we added 10 mL of DMEM supplemented with 10% FBS to neutralize the trypsin, and collected the cell suspension into a 15 mL tube. We centrifuged the samples at 200 × g for 5 minutes. After centrifugation, the resulting cell pellet was resuspended in 1 mL of PBS and transferred to a 1.5 mL Eppendorf tube. We centrifuged the suspension again at 200 × g for 5 minutes. We discarded supernatant and stored the final cell pellet at -80°C until shipment to the sequencing facility. There, we generated libraries using the SQK-LSK110 Ligation Sequencing kit for library preparation and long-read sequenced these using PromethION (Oxford Nanopore Technologies).

#### Assembly generation

We concatenated all generated fastq files and run Filtlong (v0.2.0)^69^ to keep reads with quality above or equal to 7 and minimum length of 5kbp. Then, we assembled ther reads with Flye (v.2.8.3)^70^ using the --nano-raw mode recommended for uncorrected ONT data and -i 2 to run internal polishing iterations. We polished the final draft Flye assembly using two rounds of Racon (v1.4.21)^71^ and one round of Medaka (v1.4.1, medaka_consensus)^72^. We purged the resulting assembly of overlaps based on read depth using Purge_dups (v1.2.5)^73^. After each step and at the end of the pipeline we assessed the genetic completeness and continuity of the assembly to evaluate intermediate values using BUSCO^74^ and assembly-stats^75^.

#### Annotation

We annotated repeats present in the genome assembly with RepeatMasker (v4.1.2)^76^ using the custom repeat library available for human. Moreover, we generated a new repeat library specific for our assembly with RepeatModeler (v1.0.11). After excluding those repeats that were part of repetitive protein families (performing a BLAST^77^ search against Uniprot^78^) from the resulting library, we run RepeatMasker again with this new library to annotate the specific repeats. We obtained the gene annotation of the *A. hybridus* genome assembly by combining transcript alignments, protein alignments and *ab initio* gene predictions.

Firstly, we retrieved a RNAseq dataset from *Ateles fusciceps* placental tissue from NCBI (accession number SRR3222426) and aligned it to the genome using STAR (v2.7.10)^79^ and minimap2 (v2.24)^80^ (splice option). We subsequently generated transcript models using Stringtie (v2.2.1)^81^ and merged these using TACO (v0.7.3)^82^. We obtained high-quality junctions to be used during the annotation process by running ESPRESSO (v1.3.0)^83^ after mapping with STAR. Finally, we produced assemblies with PASA (v2.5.2)^83,84^. To detect coding regions in the transcripts we run the TransDecoder program in the PASA package. Secondly, we downloaded the complete proteomes of human and *Ateles fusciceps* from Uniprot and aligned it to the genome using miniprot (v0.6)^85^. We performed *ab initio* gene predictions on the repeat-masked assembly with three different programs: GeneID (v1.4)^86^, Augustus (v3.5.0)^87^ and Genemark-ET (v4.71)^88^ with and without incorporating evidence from the RNAseq data. We run the gene predictions using trained parameters for human genomes, except for Genemark, which runs in a self-trained mode. Finally, we combined all the data into consensus CDS models using EvidenceModeler (v1.1.1)^84^. Additionally, we annotated UTRs and alternative splicing forms via two rounds of PASA annotation updates. We performed the functional annotation using the annotated proteins in the online server of Pannzer^89^.

#### Human orthologs retrieval

To match the results of our annotation with human genes, the program Orthofinder (v2.5.5)^90^ was run between the human Ensembl annotation and the *Ateles hybridus* annotation generated by the steps described before. For this search only one protein product per gene was used.

### Re-sequencing data analysis

#### Sample collection and processing

We analyzed 58 samples from eight species in the *Ateles* genus and two samples from the northern muriqui (*Brachyteles hypoxanthus*). This dataset included samples from captive and wild populations: wild count (*Ateles belzebuth*: 6*, A. chamek_NW*: 19*, A. chamek_SE*: 10*, A. fusciceps*: 1*, A. geoffroyi*: 1*, A. marginatus*: 1*, A. paniscus*: 3, *Brachyteles hypoxanthus*: 2), captive count (*A. fusciceps*: 13, *A. hybridus*: 2) (SData 1 meta*). Samples assigned to the *A. chamek* species were divided into two independent populations based on their sampling location in relation to the Madeira River following experts’ observations^5^ (*A. chamek_NW* and *A. chamek_SE*). The genomic data for 16 samples in the dataset was already published in Kuderna *et al.* (2023)^26^ with study accession PRJEB59576.

Three sample types were included in this project, from the most abundant to the least: blood (29), tissue (27), DNA extraction (1) and FTA-card (1). The first two sample types from wild individuals were extracted from Brazilian zoological collections in Instituto Nacional de Pesquisas da Amazônia (INPA), Universidade Federal do Amazonas (UFAM) and Instituto de Desenvolvimento Sustentável Mamirauá (IDSM). All samples were originally collected in compliance with all relevant ethical regulations. On the other hand, blood samples from captive individuals were extracted during routine veterinary checkups. Lastly, the only *A. chamek_NW* FTA-card sample was retrieved in the Yagua indigenous community of Nueva Esperanza, located in the Yavarí-Mirín River basin (04°19’53’’ S; 71°57’33’’ W; UT5: 00), a geographically isolated and well-preserved forest along the border between Brazil and Peru in the Peruvian Amazon. A blood sample from wildlife was collected by subsistence hunters as part of a wildlife conservation program. Local hunters were trained to impregnate blood from the cranial or caudal cava vein of the hunted animal on either Whatman filter paper No. 3 or FTA_ cards. The research protocols for the sampling of wildlife were approved by the Peruvian Forest and Wildlife Service (N◦ 258-2019-MINAGRI-SERFOR-DGGSPFFS; 27/05/2019) and the Institutional Animal Use Ethics Committee of the Universidad Peruana Cayetano Heredia (ref. 102142; 14/05/2019). A dried blood sample on filter paper from wildlife was exported with the approval of the Peruvian Forestry and Wildlife Service (N◦ 003258/SP-Peru, N◦ 003260/SP-Peru, BB-00017 20I-Spain and BB-00018 20I-Spain).

We extracted the DNA of the two first types of high-quality samples with the MagAttract HMW DNA extraction kit (Qiagen) and we built paired-end libraries with short inserts with the KAPA HyperPrep kit (Roche) PCR-free protocol. Using NovaSeq 6000 (Illumina), we sequenced paired-end reads with 2 × 151 + 18 + 8 bp length to retrieve an average coverage of 30x in each sample. More detailed explanation of the data generation can be found in Supplementary Methods of Kuderna *et al.* (2023)^26^. From the FTA-card sample DNA was extracted using a QIAamp DNA Investigator kit (Qiagen) following the manufacturer’s protocol.

#### Mapping, trimming and filtering raw reads

For reads extracted from all types of samples, we used seqtk mergepe (v1.3)^91^ to interleave these before trimming the adapters using cutadapt (v3.4, –interleaved)^92^. Then, we mapped these reads to the newly generated long read *Ateles hybridus* assembly with bwa-mem using default settings (v0.7.17)^93^. Next, we used bbmarkduplicates from the biobambam toolkit (v2.0.182)^94^ to mark duplicates and AddOrReplaceReadgroups (default settings) from PicardTools (v2.8.2)^95^ to add read groups to the resulting mapping files. We then discarded secondary alignments, unmapped reads and mappings with lower quality than 30 using the samtools view -F 260 -q 30 command (v1.14)^96^. Lastly, we further filtered reads with insert sizes smaller than 30 bp with a custom script for the FTA-card sample. At this stage, we calculated the mean and median depth in all samples independently of their origin with MOSDEPTH (v0.3.3)^97^ and those with mean coverage above 5x were kept.

#### Variant calling

We used the GATK toolkit (v4.1.7.0)^98^ to call variants on the generated CRAM files. To parallelize the three steps necessary to retrieve the variants, we created eighty-seven equal-size windows from the reference assembly. We used the HapplotypeCaller algorithm on each sample and genome window independently with the -ERC BP_RESOLUTION parameter. Next, all the variants in one genomic window for all samples were combined with CombineGVCFs under default settings. Final genotyping was approached jointly for all samples in a given window using GenotypeGVCFs (default settings). Out of these variants, we kept only SNPs that were biallelic using vcfbilallelic from vcflib (v10.5)^99^, and with a coverage per sample between 62 (twice median maximum depth across samples) and 5 (one third of the median minimum depth across samples). Using GATK VariantFiltration and based on GATK’s Best Practices Protocol, we excluded those SNPs not comprised by the expression “QD < 2 | FS > 60 | MQ < 40 | SOR > 3 | ReadPosRankSum < –8.0 | MQRankSum < –12.5”. Next, we merged the filtered SNPs in all genomic windows into a final VCF, where we further excluded variants with allele imbalance with frequencies outside the range of 0.25-0.75. Lastly, we excluded regions in scaffolds shorter than 0.5Mb while keeping the 118 longest ones covering 98% of the original assembly.

#### Relatedness

We used NGSRelate2 with default parameters (v2.0)^100^ to estimate the kinship coefficient (based on the *theta* parameter) among the individuals in the dataset independently by population using the Jacquard coefficients. The sample with the lowest coverage in each given pair of relatives was removed from the dataset at *theta* ≥ 0. 15.

### Decipher population structure

#### Principal component analysis

To investigate the population structure in the populations, four principal component analyses were conducted using smartPCA from EIGENSOFT (v7.2.1)^101^ on independent datasets differentiated by the unrelated individuals comprised: Full dataset, only *Ateles* individuals, individuals from populations found in Central and northern South America (*A. geoffroyi, A. fusciceps* and *A. hybridus* – trans-Andean) and individuals from those below the northern limit of the Amazon rainforest (*A. belzebuth, A. chamek_SE, A. chamek_NW* and *A. marginatus* – cis-Andean). Each of these datasets were independently filtered to remove variants with missingness above 40% (--geno 0.4) and to retain independent sites by accounting for linkage disequilibrium using default settings (--indep-pairwise 50 5 0.5) in PLINK (v1.9) ^102,103^. Principal components one and two were plotted in each case using the ggplot2^104^ package in R (v4.2.2)^105^.

#### Ancestry components estimation

We further inspected the populations’ structure examining ancestry components in the individuals. Using the full dataset with the same filters as in the PCA, ADMIXTURE software^106^ was run on *Ateles* and *Brachyteles* individuals to test the number of ancestral populations (K) ranging from 2 to 12 with 20 replicates each. The outputs were plotted in R (v4.2.2) using the ggplot2 package.

#### Phylogenetic analysis

To assess the phylogenetic relationships between the individuals in our dataset, we generated a phylogenetic tree comprising the callable genome. A bed file with 2554 windows of 1Mb was created based on the coordinates of the 118 longest scaffolds in the *Ateles hybridus* reference genome using bedtools makewindows (v2.30.0) ^107^. Considering these regions, we generated subset CRAM files per sample with samtools view (v1.14)^96^. Then, we extracted the consensus sequence from each of the mapping files using ANGSD (v0.931, -doFasta 1)^108^, setting an “N” when a position was unknown. Since all samples were mapped to the same reference genome and the same coordinates were extracted into ungapped sequences, we directly considered each multifasta of all individuals’ sequences in one genomic window to be an Mcis-Andean following ^27^. 2518 windows presented sequence data in all cases, covering 98.5% of the total reference genome. These were then independently trimmed with trimAL (v1.4.1)^109^ and used to generate an independent maximum likelihood tree in each window with iqTree 2 (v2.1.2, -B 1000)^110^ using an automatic model selection and 1000 bootstrap replicates. Finally, we employed ASTRAL (v5.7.8, default parameters)^111^ to build a multispecies coalescent tree from summarizing the 2518 1Mb-based trees. Finally, the phylogenetic tree was visualized with the interactive Tree Of Life online tool^112^.

### Exploration of genetic diversity

#### Genome wide heterozygosity

We estimated the genetic diversity of the individuals in the dataset using the genome wide heterozygosity, specifically calculating the proportion of heterozygotes in the callable base pairs of 100kbp sliding windows. Plots were generated to depict the distribution of genetic diversity genome-wide as well as the mean values in the callable genome as well as in the coding regions employing R (v4.2.2) and the ggplot2 package. We then generated a plot including a linear regression between sample coverage (x) and the calculated genome-wide heterozygosities (y) using geom_smooth(method = ”lm”). With this we wanted to identify potential biases caused by sample coverage (See Supplementary Text and Figures, “Coverage correlation to sample homozygosity and heterozygosity” section). We excluded PD_1569, PD_1570 and PD_1571 from this quantification after evaluation, given these presented abnormally high coverage and heterozygosity values in *A. fusciceps*’ distributions (Fig. S11, S12).

### Genome wide homozygosity

#### Genome wide inbreeding

We employed NGSRelate2 with default parameters (v2.0)^100^ to estimate the inbreeding coefficients of the individuals in the dataset by population using the Jacquard coefficients. Plots were generated using the ggplot2 package in R (v4.2.2). R was also used to perform a non-parametric Wilcoxon test to assess the significance of the difference between the observations in captive vs. wild individuals. We then generated a plot including a linear regression between sample coverage (x) and the calculated genome-wide inbreeding coefficients (y) using geom_smooth(method = ”lm”). With this we evaluated potential biases caused by sample coverage (See Supplementary Data and Figures, “Coverage correlation to sample homozygosity and heterozygosity” section). We did not exclude any sample after evaluating this (Fig. S13).

#### Runs of homozygosity (ROHs)

In order to identify homozygous tracts in the genomes of the sampled individuals, we used BCFtools roh (v1.14, -G 30)^96^. Runs of homozygosity (ROHs) longer than 0.5Mb were considered for further analysis. R (v4.2.2) was used to calculate the proportion of the genome in ROH (fROH) per sample and average in populations as well as to generate plots together with the ggplot2 package. We then generated a plot including a linear regression between sample coverage (x) and the calculated fROH proportions (y) using geom_smooth(method = ”lm”). With this we evaluated potential biases caused by sample coverage (See Supplementary Data and Figures, “Coverage correlation to sample homozygosity and heterozygosity” section). We did not exclude any sample after evaluating this (Fig. S14).

#### Mutational load

Given the lack of genetic information on the Ateles and Brachyteles hypoxanthus populations, to study the deleterious variation in these, the human state at coding positions had to be considered as the reference (REF) allele. To conduct this and be able to work with human coordinates, we generated a liftover chain from *Ateles hybridus* to the hg38 human genome. First, we created a .paf file using minimap2 (v2.24, -cx asm5)^113^, which we then converted to .chain file using transanno (v.0.4.5*) ^114^*. We then lifted the hard filtered *Ateles hybridus* VCF to human coordinates using Picard Tools LiftOverVCF (v2.25.4). We acknowledge this tool discards those positions where the REF allele differs from human’s, hence losing a high proportion of the variants given its distance to *Ateles hybridus*. Using SIFT^115^ we predicted the respective impact on protein function of the remaining set of variants categorizing these into synonymous, STOP, NS-tolerated and NS-deleterious mutations at the individual level.

#### Deleterious mutations enrichment in ROHs

We wanted to evaluated the proportion of *NS* − *del* mutations inside and outside runs of homozygosity as a proxy to understand how strongly a population has been able to purge these, following the hypothesis introduced in Szpiech et al. (2013)^116^. For this, the output of SIFT in human coordinates was lifted back to *Ateles hybridus’,* first using the chainSwap tool from UCSC Tools ^117^ to reverse the liftover chain and then UCSC Tools’ liftOver to retrieve the SIFT output in *Ateles hybridus* coordinates. Next, the coordinates of the identified *NS* − *del* mutations were intersected with ROH annotated coordinates taking into consideration the size category of these using BCFtools coverage by individual. We also followed these steps for the identified synonymous mutations. To obtain the reported 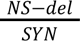 ratios, we divided the number of observed by the number of synonymous mutations inside ROHs and outside these. We divided these values with the aim of normalizing the observations by an individual baseline. Finally, we used a paired t-test (significance at *p* − *value* ≤ 0. 05) to compare the mean 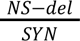 ratio per population inside and outside ROHs using R (v4.2.2).

#### Population differentiation and divergence scan

We estimated the *Fst* statistic as a proxy for population differentiation between each pair of populations throughout the genome using the genomics_general toolkit^118^. We first processed the hard filtered VCF of SNPs with all individuals using the parseVCF.py program (--skipIndels --minQual 34 --gtf flag=DP min=5 max=50). Then, we run the popgenWindows.py program specifying all populations (-w 1000 -m 200 --windType sites) to retrieve window-based statistics, including *Fst*. Subsequently, we used R (v4.2.2) to analyze the output and identify the regions in the genome driving the differentiation between *Ateles* and *Brachyteles hypoxanthus.* To do so, we kept windows in the genome that presented an *Fst* equal or larger than the 97th quantile of each population’s to *B. hypoxanthus* distribution of values and which were identified in all pairs. After that, keeping unique base pairs, we identified the span of these that intersected with the annotated coding regions in our reference genome using bedtools intersect and merge^107^ and retrieved the human orthologs of the found genes. We evaluated the functional relevance of these genes by running an overrepresentation analysis with FUMA^43^ and WebGestalt^44^, by keeping significant results (*FDR* < 0. 05) and by looking for the given SNPs in the GWAS Catalog^119^ without any relevant detection.

### Migration and gene flow

#### Deviations from isolation by distance

To investigate the potential role of the Madeira River as an effective barrier to the two hypothesized populations of *A. chamek*, we estimated deviations from Isolation by distance on 30 individuals from this species with the EEMS^120^ software. For this, we generated a genetic dissimilarity matrix using the bed2diffs_v1 program in the toolkit and together with the geographic locations of the sampled individuals, we used the geographic range distribution provided by the IUCN Red List of Threatened Species^9^. The runeems_snps program was run for 9M iterations with a thinning of 9999 and a grid density of 700 demes. Finally, we assessed the convergence of the MCMC chain by running the program more than once independently and plotted the output of this using the reemsplots2^121^ package in R (v4.2.2).

#### Gene flow testing

We used the DSUITE package to calculate D and *f*-branch statistics on the final VCF where first the cis-Andean group of samples were subsetted (*A. belzebuth, A. chamek_SE, A. chamek_NW* and *A. marginatus*) together with *Brachyteles hypoxanthus* as outgroup. And then, in a second independent run, *A. belzebuth* and *A. chamek_NW* samples were discarded and PD_0074, PD_1312 and PD_1314 *A. chamek_SE* were labeled as an independent population (*A. chamek_SEAR*) to examine their apparent genetic closeness to *A. marginatus* in contrast to the rest of the samples in the population. We focused only on the subset of cis-Andean individuals to avoid reference bias given the assembly species is in the dataset (*Ateles hybrius*) and these populations are equally distant from it phylogenetically. We then used the Dtrios, Fbranch and dtools.py programs to estimate the *f*-branch values and reported significant observations. Then, to examine these patterns at the genome-wide scale we run Dinvestigate in a sliding window fashion (w=1000, s=200). The output of this was examined…

## Acknowledgements

We gratefully acknowledge Prof. Ben Evans for providing useful insights to the interpretation of the data. T.M.B. gratefully acknowledges the financial support from the European Research Council (ERC) under the European Union’s Horizon 2020 research and innovation program (grant agreement no. 864203), (PID2021-126004NB-100) (MICIIN/FEDER, UE) and from the Secretaria d’Universitats i Recerca and CERCA Program del Departament d’Economia i Coneixement de la Generalitat de Catalunya (GRC 2021 SGR 00177). J.P.B. gratefully acknowledges the financial support from the Natural Environment Research Council (NERC) (NE/T000341/1). N.H.-A. gratefully acknowledges the financial support from the Government of Catalonia | Agència de Gestió d’Ajuts Universitaris i de Recerca (Agency for Management of University and Research Grants) (FI_00040). F.E.S. gratefully acknowledges the financial support from the Fonds National de la Recherche Scientifique (F.R.S.-FNRS, Belgium; grant 40017464), Brazilian National Council for Scientific and Technological Development (CNPq) (Processes 303286/2014-8, 303579/2014-5, 200502/2015-8, 302140/2020-4, 300365/2021-7, 301407/2021-5, #301925/2021-6), the International Primatological Society (Conservation grant), the Margot Marsh Biodiversity Foundation (SMA-CCO-G0023, SMA-CCOG0037), the Primate Conservation Inc. (1713 and 1689) and the Gordon and Betty Moore Foundation (Grant 5344) (Mamirauá Institute for Sustainable Development).

