## Supplementary Text and Figures for "The genomic landscape of spider monkeys and northern muriquis from a conservation perspective"

#### Supplementary data and figures

#### Table of Contents

**Supplementary Results** 2
- Phylogenetics 2
- Principal component analysis 3
- Ancestry components estimation 6
- Runs of homozygosity 8
- Genome wide heterozygosity 9
- Coverage correlation to sample homozygosity and heterozygosity 11
  - Genome-wide heterozygosity 11
  - Genome-wide inbreeding coefficients 14
  - Runs of homozygosity 15
- Mutational load 16
  - Deleteriousness overlap with runs of homozygosity 18
- Gene flow estimation 20

**Supplementary References** 22

### Supplementary Results

#### Phylogenetics

ASTRAL multispecies phylogenetic tree summarizing 2518 independent maximum likelihood gene trees from 1Mb genomic windows.

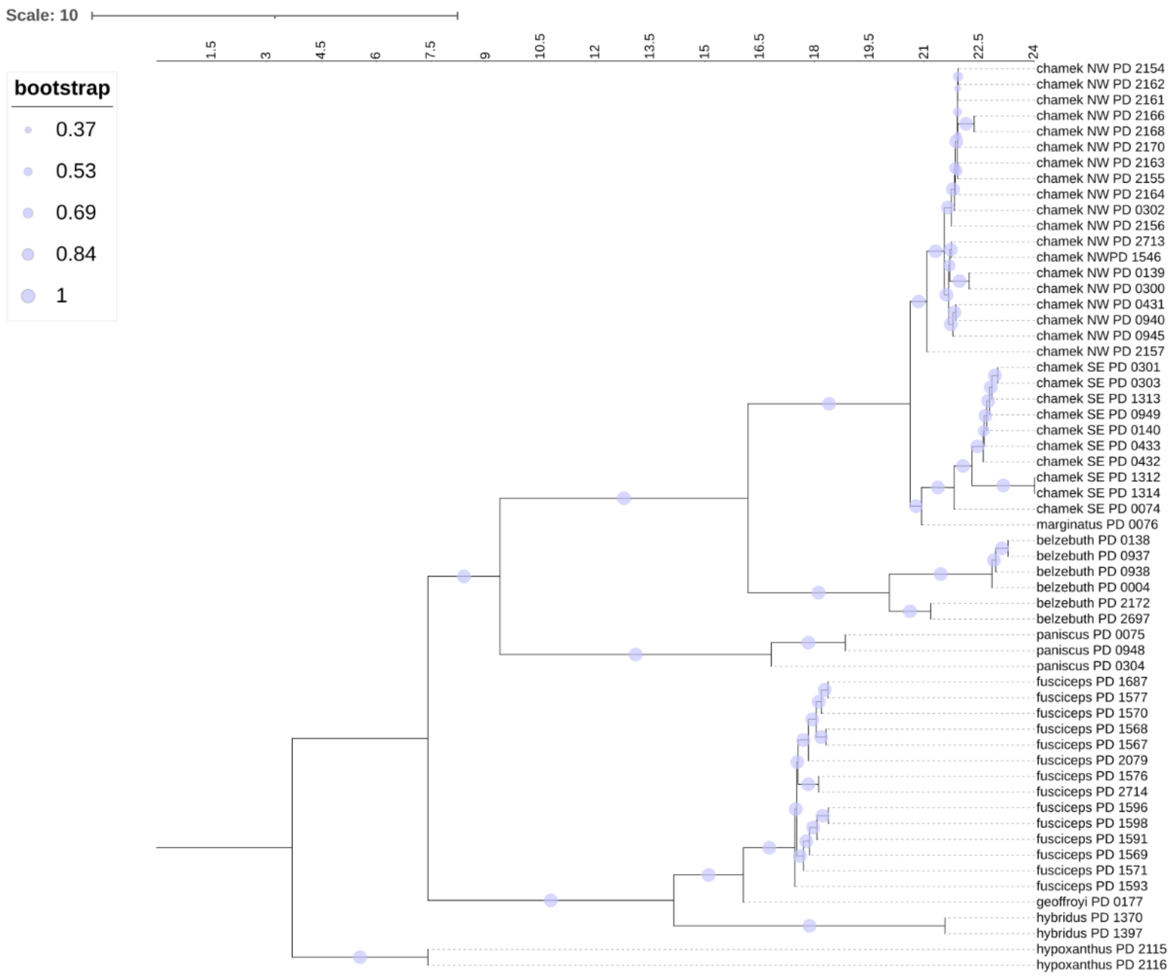


**Fig. S1. Genome-wide multispecies phylogenetic tree**. Branch lengths are proportional to coalescent units (ASTRAL) and bootstrap support is indicated by circle size.

#### Principal component analysis

PCA on distinct subsets of samples in the dataset using smartPCA from EIGENSOFT to further explore population structure.

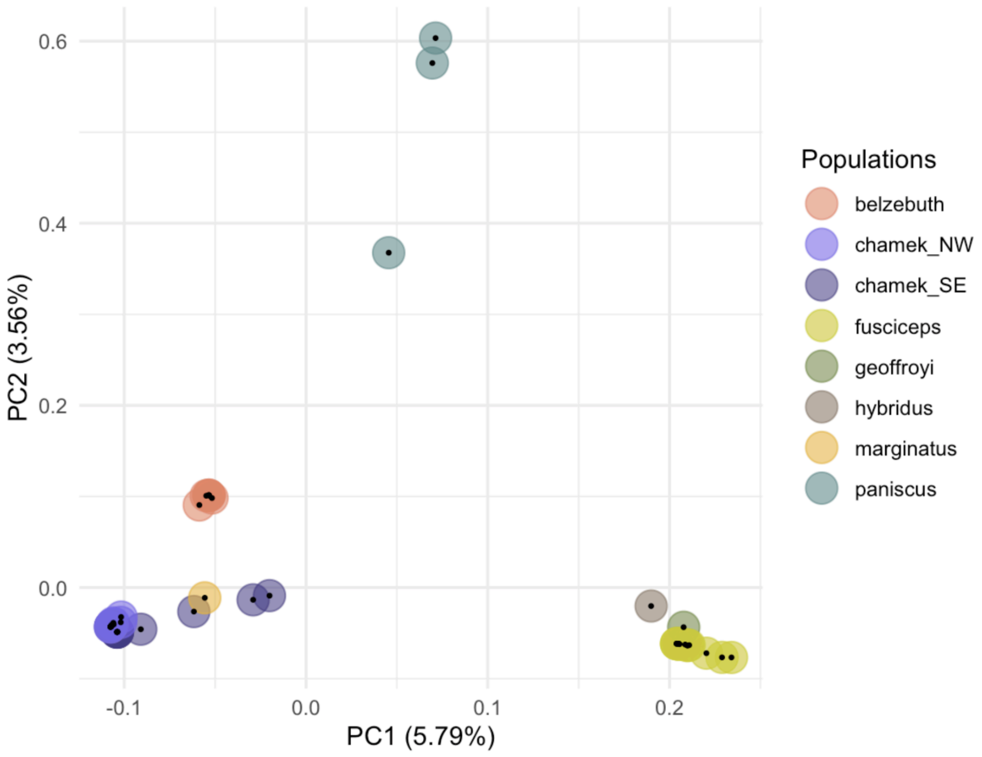


**Fig. S2. Principal component analysis of all *Ateles* unrelated samples**. Whole genome set of SNPs filtered to remove variants with missingness above 40% (--geno 0.4) and to retain independent sites by accounting for linkage disequilibrium using default settings (--indep-pairwise 50 5 0.5) yielding 55.6M SNPs.

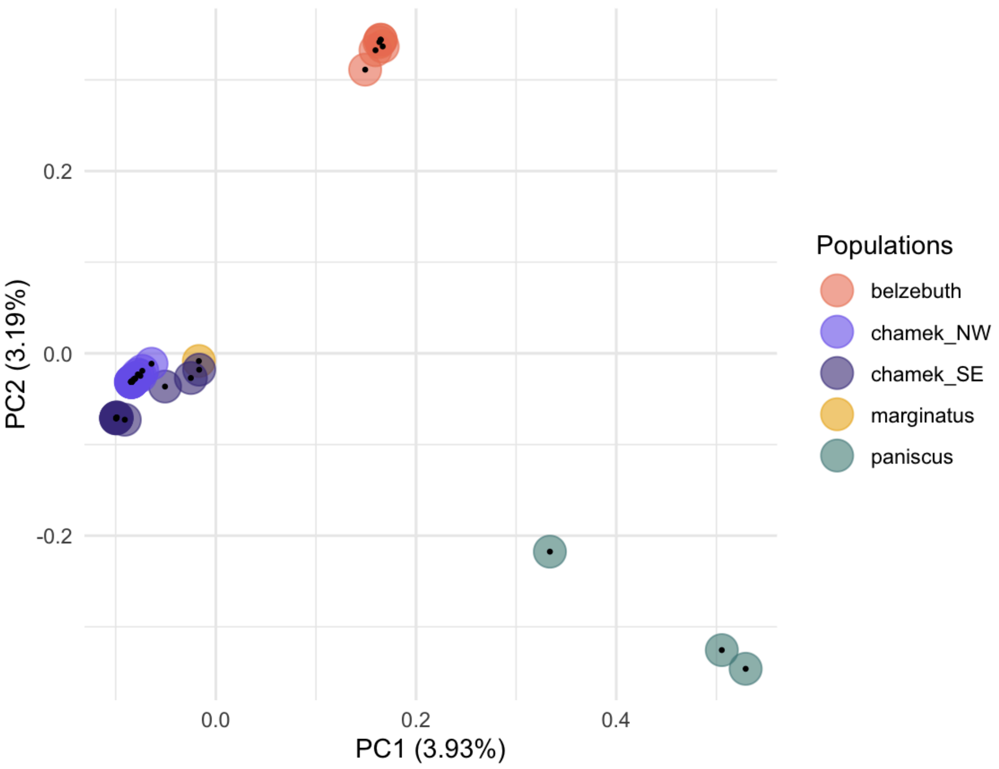


**Fig. S3. Principal component analysis of unrelated samples from amazonian *Ateles*.** Whole genome set of SNPs filtered to remove variants with missingness above 40% (--geno 0.4) and to retain independent sites by accounting for linkage disequilibrium using default settings (--indep-pairwise 50 5 0.5) yielding 21.6M SNPs.

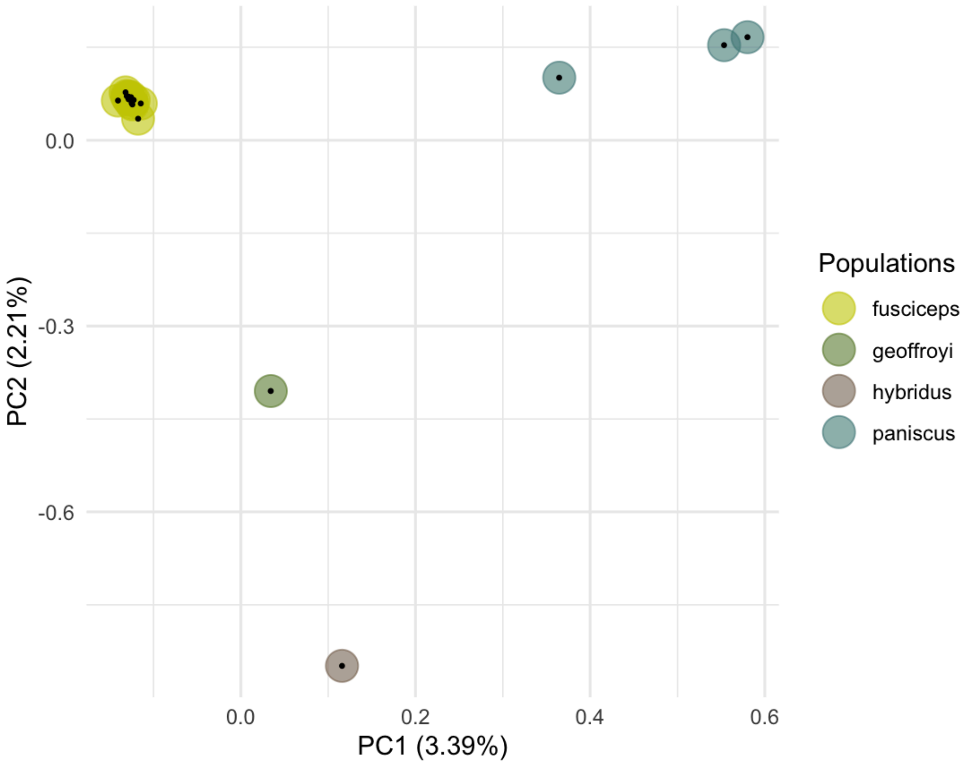


**Fig. S4. Principal component analysis of unrelated samples from non-amazonian *Ateles* and *A. paniscus***. Whole genome set of SNPs filtered to remove variants with missingness above 40% (--geno 0.4) and to retain independent sites by accounting for linkage disequilibrium using default settings (--indep-pairwise 50 5 0.5) yielding 21.6M SNPs.

#### Ancestry components estimation

Best K depicts the following population structuring, which mirrors actual results backed up from other analyses in some aspects, but is not truthful in other cases. In particular, this is not optimal for species with very low numbers of samples, then, we hypothesize biased results from other observations are due to the unevenness of samples per population. We also show $K$=8, which though exhibiting a high cross-validation error, it is reported as a reflection of population structure rather than actual ancestry sharing, in particular interest for *A. marginatus* and *A. chamek_SE* samples in the interfluvial region of the Aripuanã and Tapajos rivers .

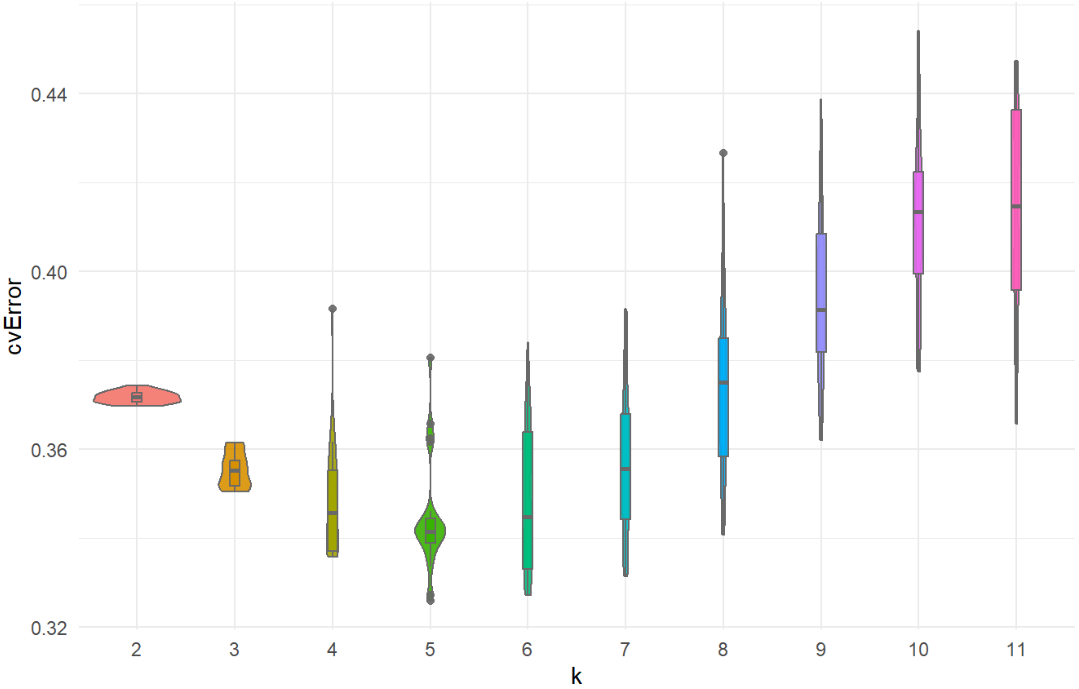


**Fig. S5. Cross validation error in ADMIXTURE**. Cross validation error calculated in each of 20 replicates per K on the whole dataset of unrelated samples with the same filters specified above in PCA. Lowest cross validation error depicts best $K$ or number of ancestral populations, here $K$=5.

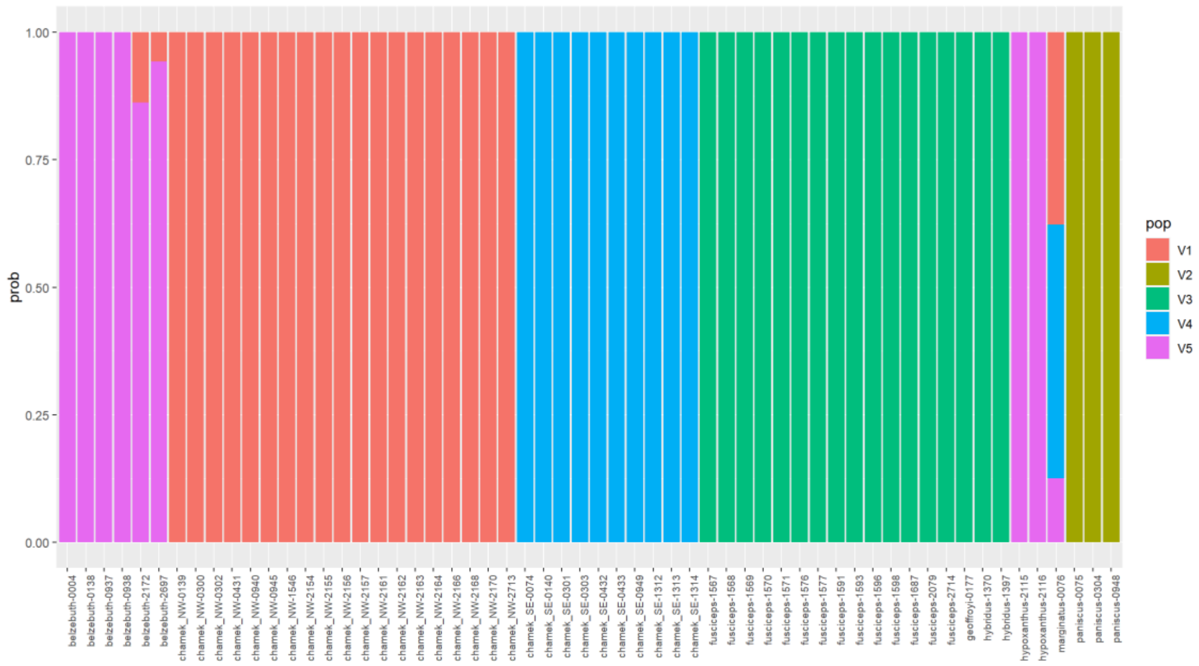


**Fig. S6. ADMIXTURE ancestry sharing proportions at best $K$=5.**

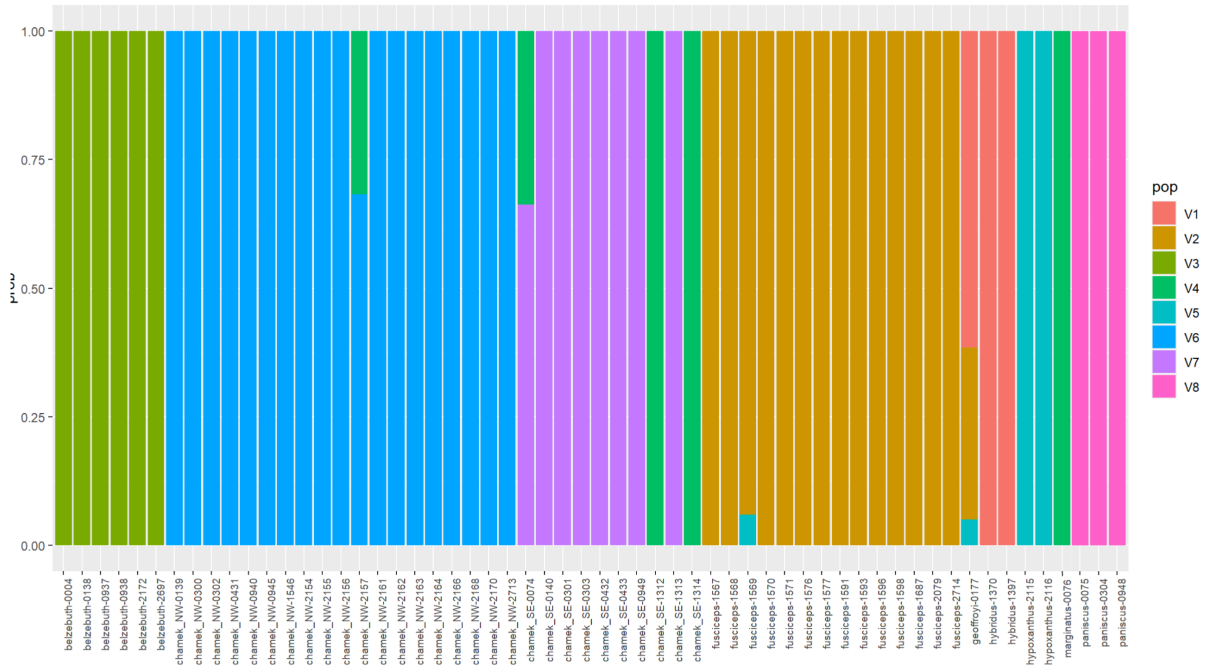


**Fig. S7. ADMIXTURE ancestry sharing proportions at $K$=8.**

#### Runs of homozygosity

The average counts per population were reported in the main text, yet variation is found in particular in some of these, notably for *A. fusciceps* managed individuals and *A. chamek_SE*.

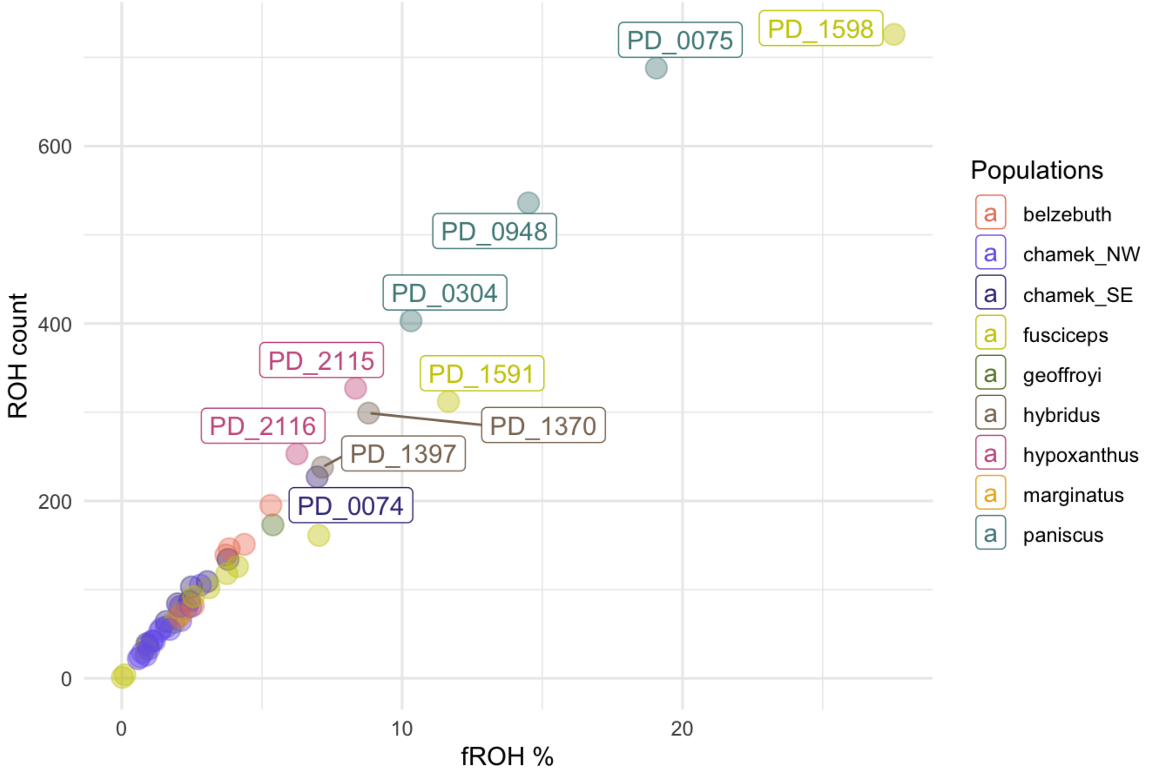


**Fig. S8. ROH count and fROH correlation.** Count of runs of homozygosity above 0.5Mb per individual correlated with the percentage of the genome in it in ROH or fROH.

#### Genome wide heterozygosity

These violin plots evidence the lack of overlap between the genome-wide heterozygosity distributions between *A. chamek_SE* and *A. chamek_NW,* giving support to their independent examination.

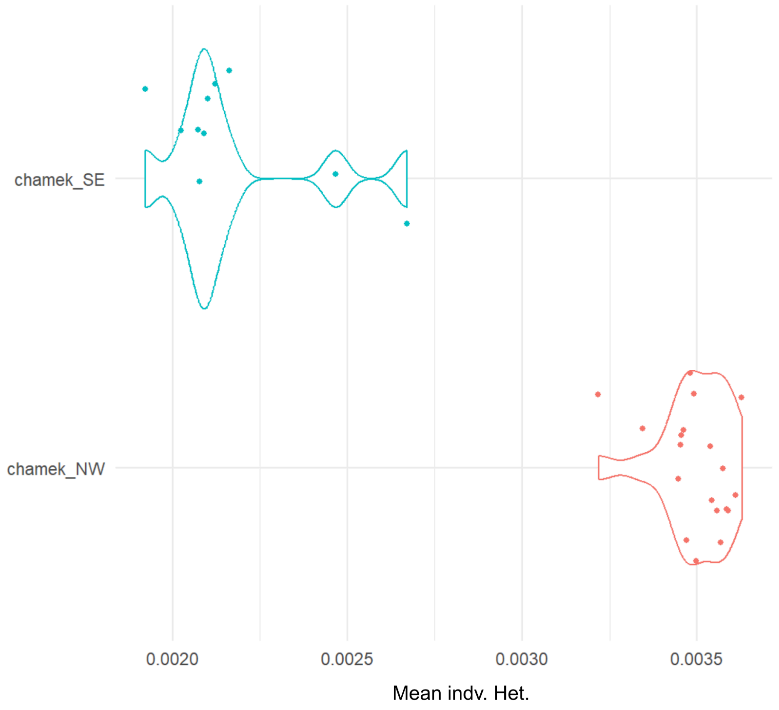


**Fig. S9. Genome-wide heterozygosity values of *A. chamek* individuals.** Samples clustered by its sampling location with respect to the Madeira River (Fig. 3).

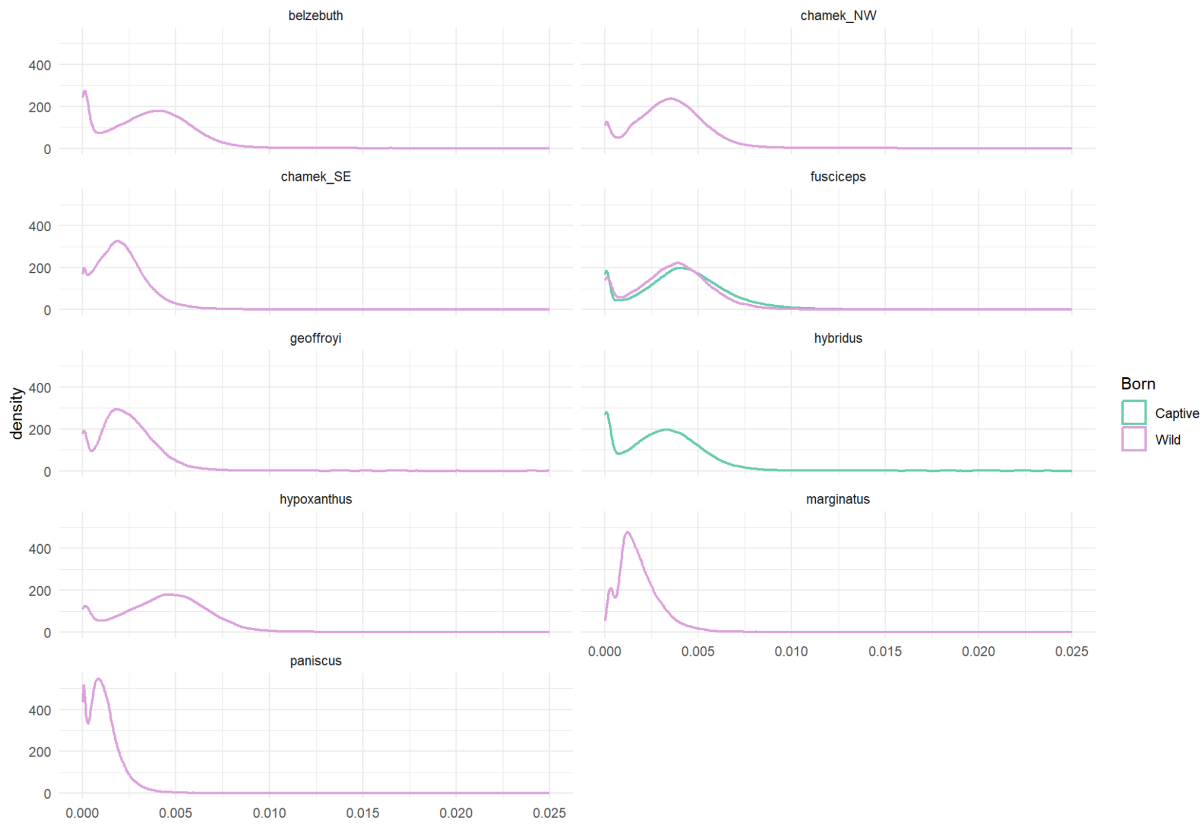


**Fig. S10. Genome-wide heterozygosity values distribution by population.** Genome-wide counts of heterozygous positions in 50kb windows summarized by population and coloured by quality of captive managed or wild.

#### Coverage correlation to sample homozygosity and heterozygosity

In our dataset some of the samples can vary substantially in terms of coverage. Low coverage has been reported to affect estimates of genome-wide heterozygosity as well as those of inbreeding (genome-wide and the detection of runs of homozygosity) [1]. The degree to which this can impact these estimates may also depend on other factors such as unidentified population substructure, which has already been addressed here.

In order to assess the potential impact of coverage in our estimates and given the properties of our dataset, we tested the correlation between coverage and several parameters linked to the homozygosity and heterozygosity of the samples. Based on this, we finally excluded only some samples for heterozygosity estimation, where we identified a relevant effect of coverage as to avoid biased conclusions. The criteria used were the following: in the identification of the proportion of heterozygous positions through the genome, we excluded samples showing a positive correlation with coverage that yielded estimates that surpassed another population's range and implied a 2-fold increase or more from samples with non-abnormally high coverage. We followed the opposite reasoning now focusing on negative correlations and decreases in homozygosity-related estimates. We were unable to assess potential biases on populations with one sample or two samples with very similar coverage.

##### Genome-wide heterozygosity

The linear model below evidenced how the identification of heterozygous positions is positively correlated with coverage in *A. fusciceps*. We excluded PD_1569, PD_1570 and PD_1571 from this quantification since these presented abnormally high coverage and heterozygosity values in *A. fusciceps*' distributions. Though we observed high uncertainty the estimated regression line, these samples' estimates cause the maximum heterozygosity in this species to increase more than 5-fold (from 0.0045 to 0.025).

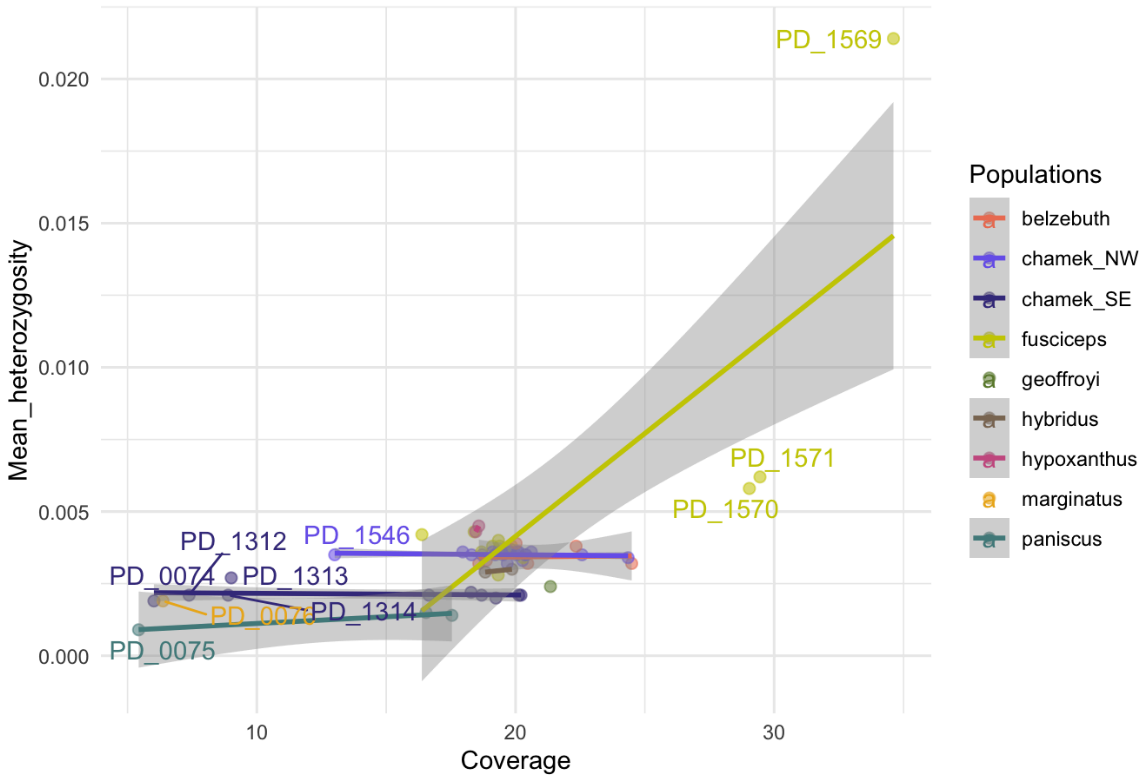


**Fig. S11. Mean heterozygosity and coverage correlation.** This plot includes all the samples in the dataset, which are represented by each dot coloured in accordance with their respective population of origin. Lines indicate regression trend and gray shade around indicates its confidence interval.

After omitting *A. fusciceps'* PD_1569, PD_1570 and PD_1571, we further identified positive correlations in *A. paniscus*, *A. hybridus* and *B. hypoxanthus*. Nevertheless, i) the variance in coverage and heterozygosity in the species was not large nor the caused increase with respect to others. Moreover, ii) given the limited number of samples, we were unable to discern between the effect of coverage from actual population variability. The latter is evidenced by *A. paniscus'* regression line's wide confidence interval.

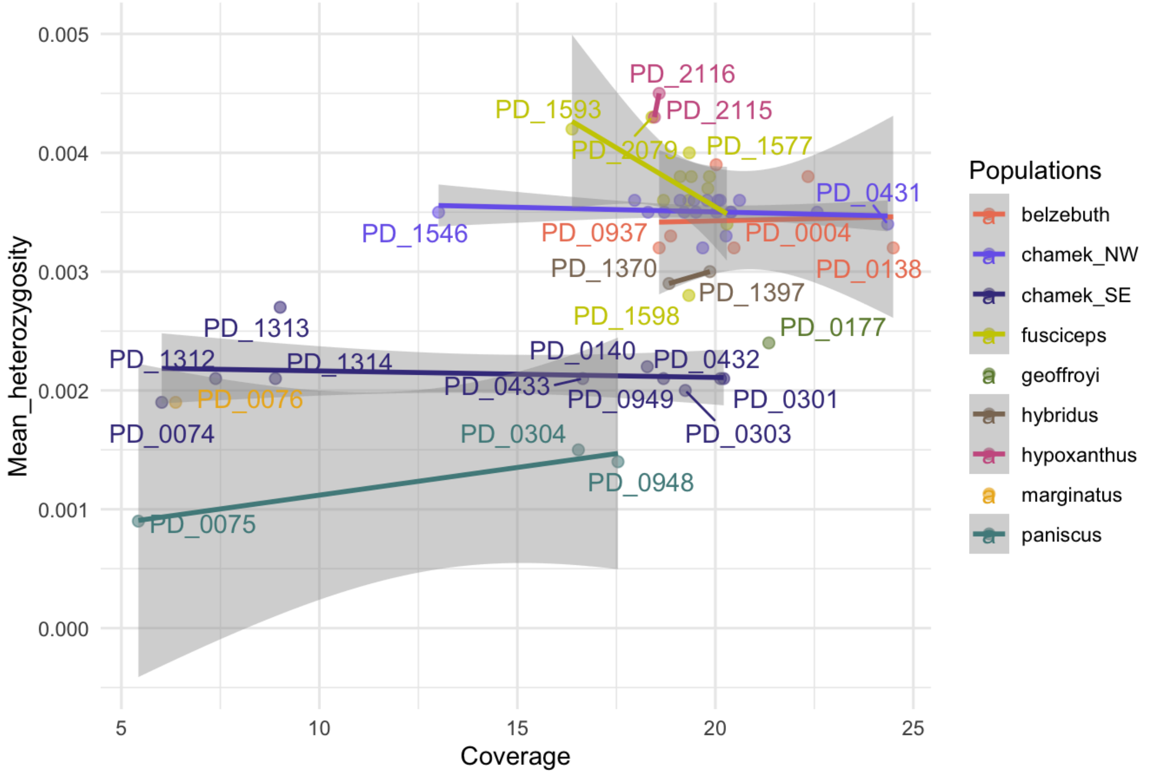


**Fig. S12. Corrected mean heterozygosity and coverage correlation.** This plot includes all the samples in the dataset but *A. fusciceps* PD_1569, PD_1570 and PD_1571, which presented abnormally high coverage values. Samples are represented by each dot coloured in accordance with their respective population of origin. Lines indicate regression trend and gray shade around indicates its confidence interval.

##### Genome-wide inbreeding coefficients

We observed a negative correlation between genome-wide inbreeding coefficients and coverage in *A. fusciceps*, *A. belzbuth* and both *A. chamek* populations. Nevertheless, here we kept all the samples in the dataset. Overall, in these populations we observed that samples of similar coverage presented great variance in genome-wide inbreeding estimates as well, ruling out a technical-driven signal. This was the case also in *A. fusciceps,* where many intermediate-coverage samples presented values of 0 in this estimate as the greatly coverage-differentiated PD_1569, PD_1570 and PD_1571. These choices were further backed up by the observed uncertainty in the regression coefficients found.

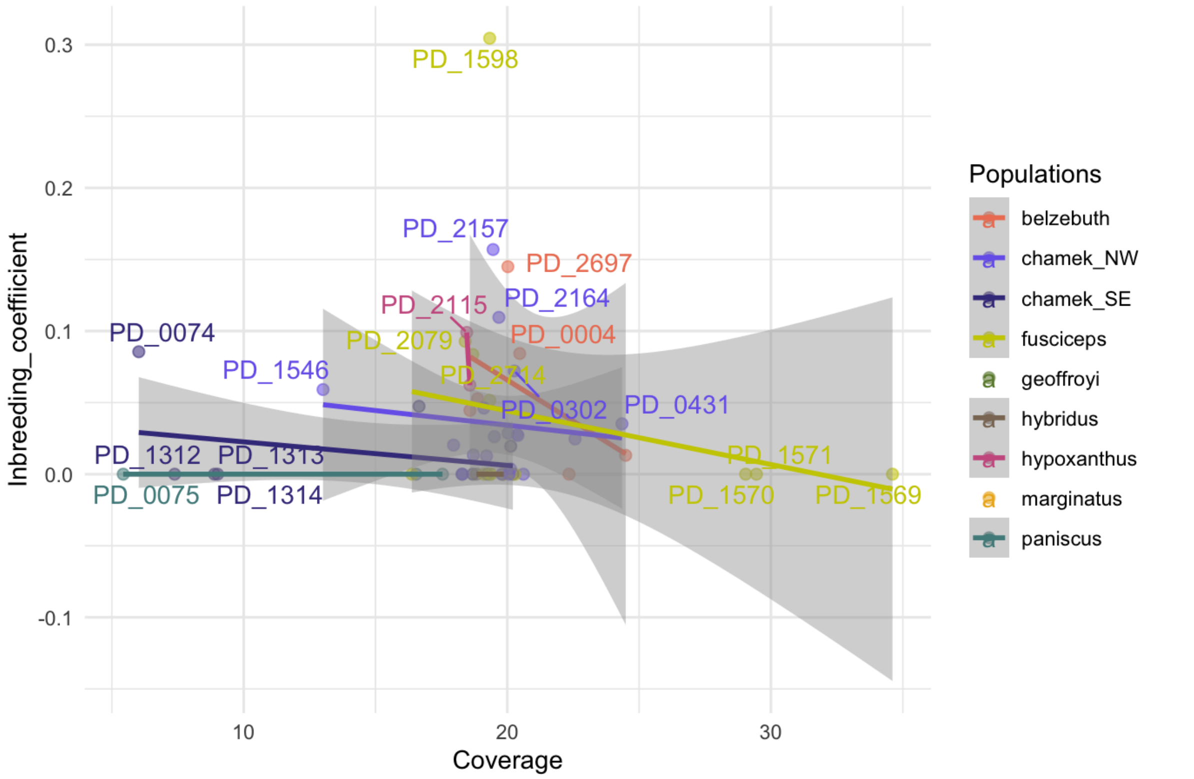


**Fig. S13. Genome-wide inbreeding coefficients and coverage correlation.** This plot includes all the samples in the dataset, which are represented by each dot coloured in accordance with their respective population of origin. Lines indicate regression trend and gray shade around indicates its confidence interval.

#### Runs of homozygosity

We did not identify any relevant contribution from coverage variation to the estimate of the total proportion of the genome in runs of homozygosity (fROH) in our dataset following the criteria introduced above.

The latter was also the case at the ROH count level. Through the joint evaluation of the observed coverage distribution and the exposed patterns in Fig. S8, we found that the identified signals were not coverage-driven.

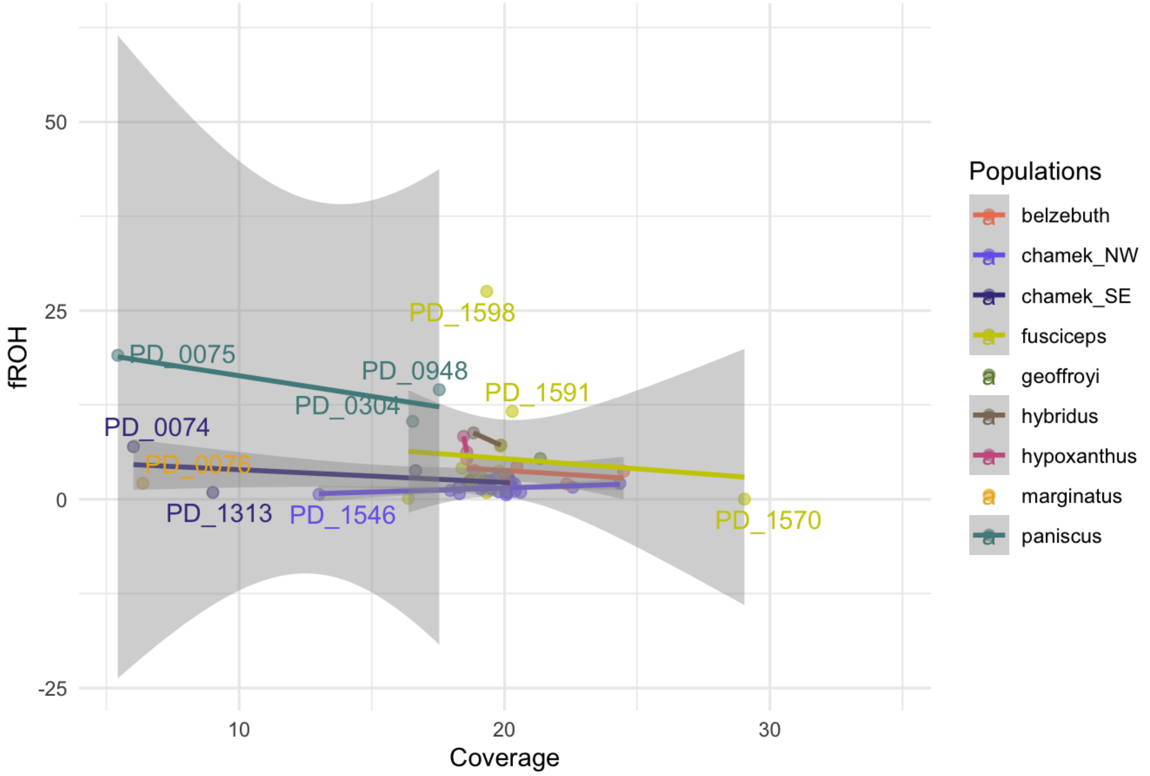


**Fig. S14. Total proportion of genome in ROH (fROH) and coverage correlation.** This plot includes all the samples in the dataset, which are represented by each dot coloured in accordance with their respective population of origin. Lines indicate regression trend and gray shade around indicates its confidence interval.

#### Mutational load

Reported proportions of genome-wide non-synonymous (NS) mutations were normalized by synonymous ones (SYN). Population-level averages were reported, though as it can be appreciated, there is intra-population variation. The potential impact of coverage variance on the results of this analysis was not explicitly assessed, as the provided estimates were normalized by the number of synonymous (SYN) mutations identified in each sample. While this normalization accounts for differences in sequencing depth, it does not eliminate the possibility that low coverage may have affected variant detection. However, we are unable to quantify the extent of this potential bias.

PD_2079 captive *A. fusciceps* stands out as well as in genome-wide inbreeding (Fig. 2B), while remaining managed individuals are spread through the distribution. On the other hand, highlighted PD_1312 and PD_1314 were sampled at the northernmost point and most isolated area of the interfluvial region of the Aripuanã and Tapajos rivers from the remaining *A. chamek_SE* distribution (Fig. 3), which can potentially be linked to the observed amounts of genetic load exhibited here as $\frac{NS-del}{SYN}$ ratios. While PD_0074 shared many properties with these two samples in terms of population dynamics and gene flow with *A. marginatus*, here it does not exhibit as strong signs of isolation, in agreement with its sampling point, which is found very close to *A. marginatus*'s distribution, but southern and hence not as disconnected from the remaining *A. chamek_SE* individuals than the other two (Fig. 3).

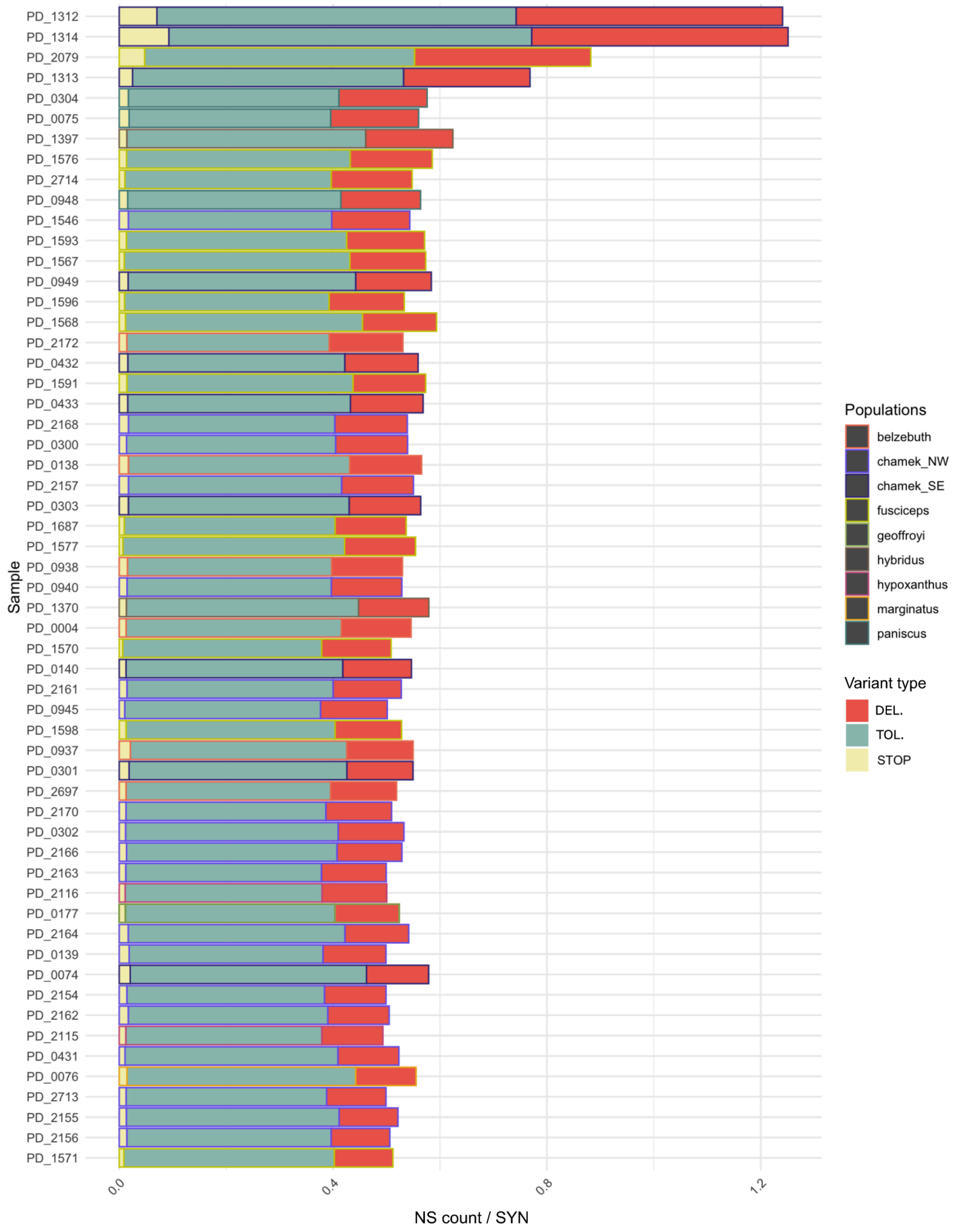


**Fig. S15. Non-synonymous (NS) variation proportions in samples.** Ratio of non-synonymous mutations by synonymous coloured by NS mutation type as predicted by SIFT: deleterious (DEL.), tolerated (TOL.) and STOP; and highlighted by population color pattern.

#### Deleteriousness overlap with runs of homozygosity

The rationale behind this test is that under normal circumstances a population will show smaller proportion of mildly non-synonymous deleterious mutations in runs of homozygosity than in the rest of the genome, since this are usually purged given their deleteriousness would be even more harmful in homozygosity [2]. Therefore, if some population has struggled to purge this mildly deleterious variation, this ratio would be increased in ROHs when compared to the rest of the genome. In this line, *A. belzebuth* and *A. chamek_SE* yielded significant results in the comparison of the mean population ratios in and outside ROHs, in the first being this higher inside ROHs and in *A. chamek_SE* outside. The ratio was also higher in *A. hybridus* but it was only close to being significant.

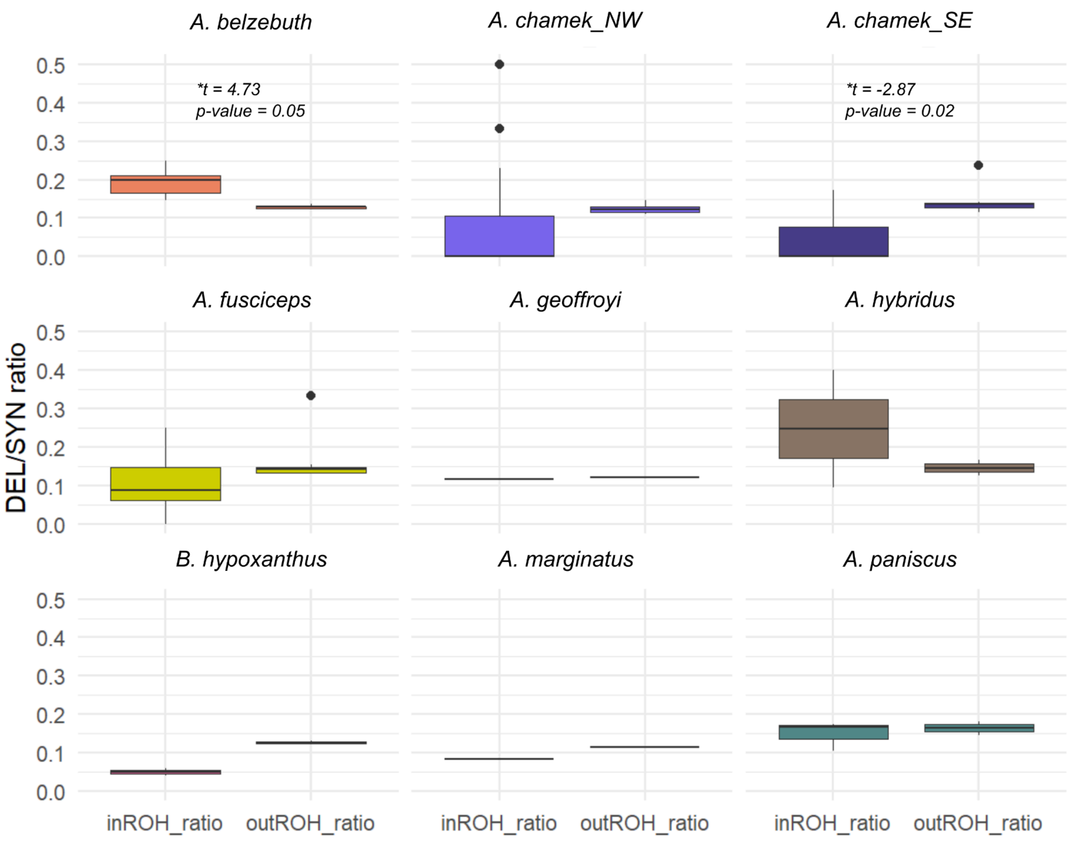


**Fig. S16. Proportion of $\frac{NS{-}del}{SYN}$ in and outside runs of homozygosity (ROHs).** Proportion of NS-deleterious mutations normalized by the count of synonymous mutations inside and outside of ROHs independently. Boxplots indicate distribution and mean values in populations. Significant comparisons at $p - value \leq 0.05$ of population mean ratios using paired *t-test* indicated in figure.

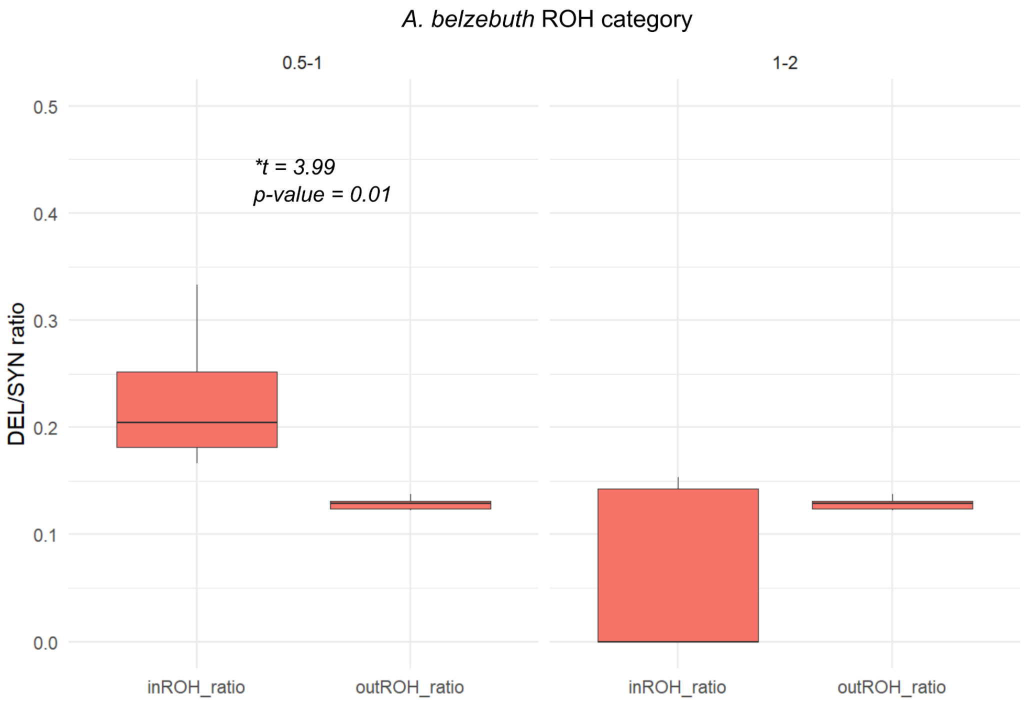


**Fig. S17. Proportion of $\frac{NS-del}{SYN}$ in and outside runs of homozygosity (ROHs) by length category in *A. belzebuth*.** Significant comparisons at $p - value \leq 0.05$ of mean ratios per ROH length category using paired *t-test* indicated in figure.

#### Gene flow estimation

Since *Ateles hybridus*, the species of the reference genome in this study was included in the dataset, we only inspected the connectivity patterns between species that were roughly equally distant from it, therefore, these analyses were based on amazonian spider monkeys with *Brachyteles hypoxanthus* as outgroup. This analysis was conducted twice, the second time focusing only on *A. chamek_NW, A. chamek_SE* and *A. marginatus* with the aim of investigating the gene flow between the last two. In line with this, PD_0074, PD_1312 and PD_1314 were treated as an independent population from the remaining *A. chamek_SE* as in other analyses these showed signs of strengthened connectivity with *A. marginatus* (Fig. 1B, S1-3, S7). When this second analysis was performed, the remaining *A. chamek_SE* individuals did not show signs of gene flow with *A. marginatus* anymore, which would not mean there hasn't been connectivity between these, but that the signal exhibited by the aforementioned three individuals is far stronger and/or much more recent than what could be detected in the other *A. chamek_SE*.

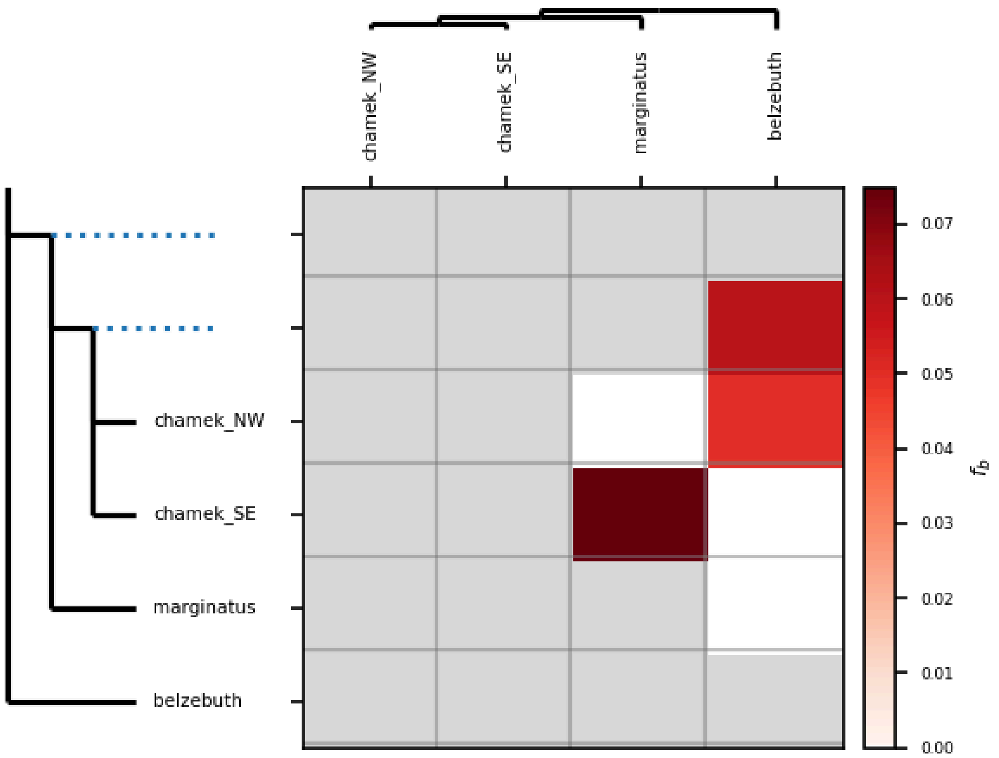


**Fig. S18. DSUITE $f$-branch values on all amazonian *Ateles*.** Significant values reported at $FDR < 0.05$ using *Brachyteles hypoxanthus* as an outgroup.

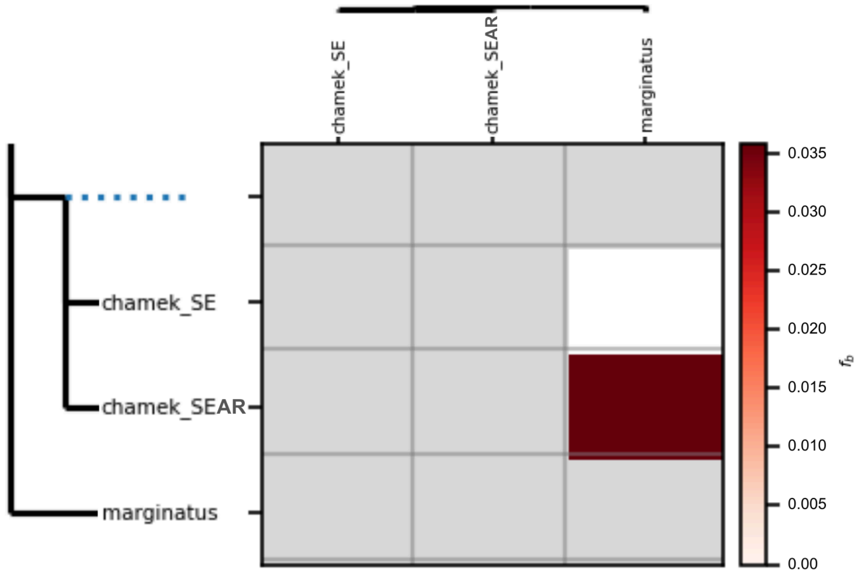


**Fig. S19. DSUITE $f$-branch values on *A. marginatus* and *A. chamek_SE* split into *A. chamek_SEAR*** (PD_0074, PD_1312 and PD_1314) **and all other individuals.** Significant values reported at $FDR < 0.05$ using *Brachyteles hypoxanthus* as an outgroup.
