## Supplementary Data 1 for "The genomic landscape of spider monkeys and northern muriquis from a conservation perspective"

| Sample_ID | Genus | Species | Population | Born | Sample_type | IUCN |
|---|---|---|---|---|---|---|
| PD_0004 | *Ateles* | *belzebuth* | *belzebuth* | Wild | Tissue | EN |
| PD_0074 | *Ateles* | *chamek* | *chamek_SE* | Wild | Tissue | EN |
| PD_0075 | *Ateles* | *paniscus* | *paniscus* | Wild | Tissue | VU |
| PD_0076 | *Ateles* | *marginatus* | *marginatus* | Wild | Tissue | EN |
| PD_0138 | *Ateles* | *belzebuth* | *belzebuth* | Wild | Tissue | EN |
| PD_0139 | *Ateles* | *chamek* | *chamek_NW* | Wild | Tissue | EN |
| PD_0140 | *Ateles* | *chamek* | *chamek_SE* | Wild | Tissue | EN |
| PD_0177 | *Ateles* | *geoffroyi* | *geoffroyi* | Wild | Tissue | EN |
| PD_0300 | *Ateles* | *chamek* | *chamek_NW* | Wild | Tissue | EN |
| PD_0301 | *Ateles* | *chamek* | *chamek_SE* | Wild | Tissue | EN |
| PD_0302 | *Ateles* | *chamek* | *chamek_NW* | Wild | Tissue | EN |
| PD_0303 | *Ateles* | *chamek* | *chamek_SE* | Wild | Tissue | EN |
| PD_0304 | *Ateles* | *paniscus* | *paniscus* | Wild | Tissue | VU |
| PD_0431 | *Ateles* | *chamek* | *chamek_NW* | Wild | Tissue | EN |
| PD_0432 | *Ateles* | *chamek* | *chamek_SE* | Wild | Tissue | EN |
| PD_0433 | *Ateles* | *chamek* | *chamek_SE* | Wild | Tissue | EN |
| PD_0937 | *Ateles* | *belzebuth* | *belzebuth* | Wild | Tissue | EN |
| PD_0938 | *Ateles* | *belzebuth* | *belzebuth* | Wild | Tissue | EN |
| PD_0940 | *Ateles* | *chamek* | *chamek_NW* | Wild | Tissue | EN |
| PD_0945 | *Ateles* | *chamek* | *chamek_NW* | Wild | Tissue | EN |
| PD_0948 | *Ateles* | *paniscus* | *paniscus* | Wild | Tissue | VU |
| PD_0949 | *Ateles* | *chamek* | *chamek_SE* | Wild | Tissue | EN |
| PD_1312 | *Ateles* | *chamek* | *chamek_SE* | Wild | Tissue | EN |
| PD_1313 | *Ateles* | *chamek* | *chamek_SE* | Wild | Tissue | EN |
| PD_1314 | *Ateles* | *chamek* | *chamek_SE* | Wild | Tissue | EN |
| PD_1370 | *Ateles* | *hybridus* | *hybridus* | Captive | Whole Blood | CR |
| PD_1397 | *Ateles* | *hybridus* | *hybridus* | Captive | Whole Blood | CR |
| PD_1546 | *Ateles* | *chamek* | *chamek_NW* | Wild | FTA-card | EN |
| PD_1567 | *Ateles* | *fusciceps* | *fusciceps* | Captive | Whole blood | EN |
| PD_1568 | *Ateles* | *fusciceps* | *fusciceps* | Captive | Whole blood | EN |
| PD_1569 | *Ateles* | *fusciceps* | *fusciceps* | Captive | Whole blood | EN |
| PD_1570 | *Ateles* | *fusciceps* | *fusciceps* | Captive | Whole blood | EN |
| PD_1571 | *Ateles* | *fusciceps* | *fusciceps* | Captive | Whole blood | EN |
| PD_1576 | *Ateles* | *fusciceps* | *fusciceps* | Captive | Whole blood | EN |
| PD_1577 | *Ateles* | *fusciceps* | *fusciceps* | Captive | Whole blood | EN |
| PD_1591 | *Ateles* | *fusciceps* | *fusciceps* | Captive | Whole blood | EN |
| PD_1593 | *Ateles* | *fusciceps* | *fusciceps* | Captive | whole blood | EN |
| PD_1596 | *Ateles* | *fusciceps* | *fusciceps* | Captive | Whole blood | EN |
| PD_1598 | *Ateles* | *fusciceps* | *fusciceps* | Captive | Whole blood | EN |
| PD_1687 | *Ateles* | *fusciceps* | *fusciceps* | Captive | Whole blood | EN |
| PD_2079 | *Ateles* | *fusciceps* | *fusciceps* | Captive | DNA | EN |
| PD_2115 | *Brachyteles* | *hypoxanthus* | *hypoxanthus* | Wild | Tissue | CR |
| PD_2116 | *Brachyteles* | *hypoxanthus* | *hypoxanthus* | Wild | Tissue | CR |
| PD_2154 | *Ateles* | *chamek* | *chamek_NW* | Wild | Blood | EN |
| PD_2155 | *Ateles* | *chamek* | *chamek_NW* | Wild | Blood | EN |
| PD_2156 | *Ateles* | *chamek* | *chamek_NW* | Wild | Blood | EN |

| | | | | | | |
|---|---|---|---|---|---|---|
| PD_2157 | *Ateles* | *chamek* | *chamek_NW* | Wild | Blood | EN |
| PD_2161 | *Ateles* | *chamek* | *chamek_NW* | Wild | Blood | EN |
| PD_2162 | *Ateles* | *chamek* | *chamek_NW* | Wild | Blood | EN |
| PD_2163 | *Ateles* | *chamek* | *chamek_NW* | Wild | Blood | EN |
| PD_2164 | *Ateles* | *chamek* | *chamek_NW* | Wild | Blood | EN |
| PD_2166 | *Ateles* | *chamek* | *chamek_NW* | Wild | Blood | EN |
| PD_2168 | *Ateles* | *chamek* | *chamek_NW* | Wild | Blood | EN |
| PD_2170 | *Ateles* | *chamek* | *chamek_NW* | Wild | Blood | EN |
| PD_2172 | *Ateles* | *belzebuth* | *belzebuth* | Wild | Blood | EN |
| PD_2697 | *Ateles* | *belzebuth* | *belzebuth* | Wild | Blood | EN |
| PD_2713 | *Ateles* | *chamek* | *chamek_NW* | Wild | Blood | EN |
| PD_2714 | *Ateles* | *fusciceps* | *fusciceps* | Wild | Blood | EN |

| Sample_ID | Coverage | Mean_heterozygosity | Inbreeding_coeffiicient |
|---|---|---|---|
| PD_0004 | 20,47 | 0,0032 | 0,084259 |
| PD_0074 | 6,02 | 0,0019 | 0,085521 |
| PD_0075 | 5,43 | 0,0009 | 0 |
| PD_0076 | 6,37 | 0,0019 | NA |
| PD_0138 | 24,49 | 0,0032 | 0,013189 |
| PD_0139 | 22,57 | 0,0035 | 0,024623 |
| PD_0140 | 18,28 | 0,0022 | 0 |
| PD_0177 | 21,35 | 0,0024 | NA |
| PD_0300 | 20,61 | 0,0036 | 0 |
| PD_0301 | 20,21 | 0,0021 | 0 |
| PD_0302 | 20,27 | 0,0033 | 0,072076 |
| PD_0303 | 19,24 | 0,0020 | 0 |
| PD_0304 | 16,54 | 0,0015 | 0 |
| PD_0431 | 24,35 | 0,0034 | 0,035175 |
| PD_0432 | 20,13 | 0,0021 | 0,019582 |
| PD_0433 | 16,65 | 0,0021 | 0,047417 |
| PD_0937 | 18,58 | 0,0032 | 0,044543 |
| PD_0938 | 18,87 | 0,0033 | 0,052619 |
| PD_0940 | 17,96 | 0,0036 | 0,02032 |
| PD_0945 | 18,71 | 0,0035 | 0,013344 |
| PD_0948 | 17,54 | 0,0014 | 0 |
| PD_0949 | 18,69 | 0,0021 | 0 |
| PD_1312 | 7,38 | 0,0021 | 0 |
| PD_1313 | 9,01 | 0,0027 | 0 |
| PD_1314 | 8,89 | 0,0021 | 0 |
| PD_1370 | 18,83 | 0,0029 | 0 |
| PD_1397 | 19,86 | 0,0030 | 0 |
| PD_1546 | 13,01 | 0,0035 | 0,059103 |
| PD_1567 | 19,39 | 0,0038 | 0 |
| PD_1568 | 19,81 | 0,0037 | 0 |
| PD_1569 | 34,61 | 0,0214 | 0 |
| PD_1570 | 29,04 | 0,0058 | 0 |
| PD_1571 | 29,45 | 0,0062 | 0 |
| PD_1576 | 19,33 | 0,0036 | 0,051607 |
| PD_1577 | 19,34 | 0,0040 | 0 |
| PD_1591 | 20,29 | 0,0034 | 0 |
| PD_1593 | 16,38 | 0,0042 | 0 |
| PD_1596 | 19,85 | 0,0038 | 0 |
| PD_1598 | 19,33 | 0,0028 | 0,304477 |
| PD_1687 | 19,11 | 0,0038 | 0 |
| PD_2079 | 18,40 | 0,0043 | 0,092783 |
| PD_2115 | 18,46 | 0,0043 | 0,099034 |
| PD_2116 | 18,58 | 0,0045 | 0,062175 |
| PD_2154 | 20,03 | 0,0035 | 0,028706 |
| PD_2155 | 20,40 | 0,0035 | 0,027028 |
| PD_2156 | 20,12 | 0,0036 | 0,003751 |

| | | | |
|---|---|---|---|
| PD_2157 | 19,46 | 0,0036 | 0,156916 |
| PD_2161 | 18,30 | 0,0035 | 0 |
| PD_2162 | 19,51 | 0,0035 | 0,026183 |
| PD_2163 | 20,38 | 0,0035 | 0,027677 |
| PD_2164 | 19,68 | 0,0032 | 0,109545 |
| PD_2166 | 20,07 | 0,0036 | 0 |
| PD_2168 | 19,79 | 0,0036 | 0 |
| PD_2170 | 19,21 | 0,0035 | 0,012709 |
| PD_2172 | 22,34 | 0,0038 | 0 |
| PD_2697 | 20,02 | 0,0039 | 0,144948 |
| PD_2713 | 19,11 | 0,0036 | 0,046172 |
| PD_2714 | 18,69 | 0,0036 | 0,083627 |

| Sample_ID | Latitude | Longitude |
| --- | --- | --- |
| PD_0004 | 0,4917 | -65,2717 |
| PD_0074 | -9,9783 | -56,0814 |
| PD_0075 | -1,4201 | -56,7206 |
| PD_0076 | -9 | -53 |
| PD_0138 | 0,8789 | -63,4494 |
| PD_0139 | -3,174 | -67,3895 |
| PD_0140 | -12,0558 | -60,6726 |
| PD_0177 | | |
| PD_0300 | -3,1735 | -67,3899 |
| PD_0301 | -12,49 | -63,52 |
| PD_0302 | -13,5182 | -60,4436 |
| PD_0303 | -12,491 | -63,5241 |
| PD_0304 | -2,0713 | -58,3751 |
| PD_0431 | -3,8335 | -67,4276 |
| PD_0432 | -7,7116 | -60,5968 |
| PD_0433 | -7,4607 | -60,6855 |
| PD_0937 | 0,8789 | -63,4494 |
| PD_0938 | 0,8789 | -63,4494 |
| PD_0940 | -3,8336 | -67,4278 |
| PD_0945 | -3,5649 | -65,969 |
| PD_0948 | -1,4201 | -56,7206 |
| PD_0949 | -13,5231 | -60,4345 |
| PD_1312 | -5 | -59 |
| PD_1313 | -8,7 | -63,86 |
| PD_1314 | -5 | -59 |
| PD_1370 | NA | NA |
| PD_1397 | NA | NA |
| PD_1546 | NA | NA |
| PD_1567 | NA | NA |
| PD_1568 | NA | NA |
| PD_1569 | NA | NA |
| PD_1570 | NA | NA |
| PD_1571 | NA | NA |
| PD_1576 | NA | NA |
| PD_1577 | NA | NA |
| PD_1591 | NA | NA |
| PD_1593 | NA | NA |
| PD_1596 | NA | NA |
| PD_1598 | NA | NA |
| PD_1687 | NA | NA |
| PD_2079 | NA | NA |
| PD_2115 | NA | NA |
| PD_2116 | NA | NA |
| PD_2154 | -12,187271 | -69,8479162 |
| PD_2155 | -12,187271 | -69,8479162 |
| PD_2156 | -12,187271 | -69,8479162 |

| | | |
|---|---|---|
| PD_2157 | -12,187271 | -69,8479162 |
| PD_2161 | -12,187271 | -69,8479162 |
| PD_2162 | -12,187271 | -69,8479162 |
| PD_2163 | -12,187271 | -69,8479162 |
| PD_2164 | -12,187271 | -69,8479162 |
| PD_2166 | -12,5784704 | -69,196915 |
| PD_2168 | -12,5784704 | -69,196915 |
| PD_2170 | -12,5784704 | -69,196915 |
| PD_2172 | NA | NA |
| PD_2697 | -12,0340533 | -76,92613 |
| PD_2713 | -4,1425841 | -73,4711282 |
| PD_2714 | NA | NA |
